## Supplement information for "High clonal diversity and spatial genetic admixture in early prostate cancer and surrounding normal tissue"

1. Supplementary Figures pg. 2
2. Supplementary Methods pg. 37
3. Supplementary Tables pg. 43
4. Supplementary Notes pg. 44
5. Supplementary References pg. 47

### 1. Supplementary Figures

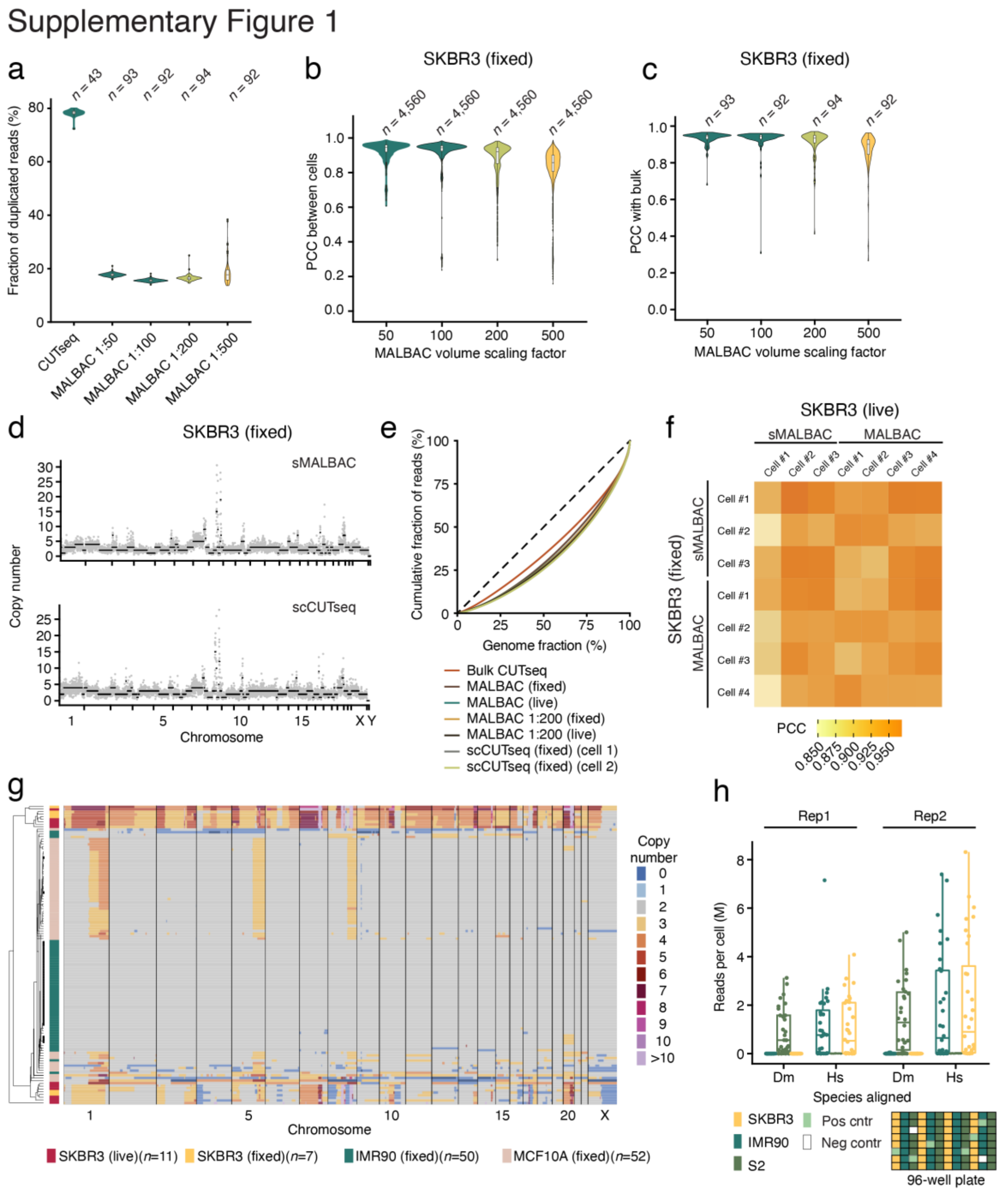


**Supplementary Fig. 1.** Technical performance and reproducibility of scCUTseq. (**a**) Fraction of read duplicates obtained after performing standard CUTseq directly on single cells or performing CUTseq after MALBAC by scaling reagent volumes 50, 100, 200 and 500 times. *n*, number of single cells with over 50K reads (**b**) Pearson's correlation coefficient (PCC) for all possible pair-wise comparisons between the segmented copy number profiles of individual SKBR3 cells processed by scCUTseq, using different MALBAC reagent volume scaling factors. *n*, number of pair-wise comparisons in each group. (**c**) Same as in (b) but comparing the copy number profile of each single cell with the copy number profile of the corresponding cell line determined by bulk CUTseq. *n*, number of cells compared to bulk CUTseq. In (a-c), violins extend from minimum to maximum, each box in the boxplot inside each violin spans from the 25th to the 75th percentile and whiskers extend from –1.5×IQR to +1.5×IQR from the closest quartile, where IQR is the inter-quartile range. Black dots, outliers. (**d**) Example of single-cell copy number profiles of fixed SKBR3 cells determined by performing a 1:200 scaled down version of MALBAC (sMALBAC) followed either by standard library preparation or by CUTseq (scCUTseq). Each gray dot represents a 500 kilobases (kb) genomic bin. Black dots indicate segmented copy number profiles determined by circular binary segmentation. (**e**) Lorenz curves for genomic coverage by bulk CUTseq, standard MALBAC, 1:200 sMALBAC, and scCUTseq on SKBR3 cells. (**f**) Correlation matrix showing the similarity between the segmented copy number profiles of individual fixed or non-fixed (live) SKBR3 cells obtained by performing standard MALBAC or 1:200 sMALBAC followed by library preparation with a commercial kit (NEBNext, see **Methods**). (**g**) Hierarchically clustered single-cell copy number profiles obtained by applying scCUTseq to four different cell lines, with or without mild fixation (4% PFA for 10 min, see **Methods**). *n*, number of single cells. (**h**) Distributions of scCUTseq reads per cell after alignment to the *Homo sapiens* (Hs) or *Drosophila melanogaster* (Dm) reference genomes for three different cell types (Hs IMR90 and SKBR3 cells; Dm S2 cells) sorted in different columns of a 96-well plate as shown in the bottom scheme. Pos cntr, positive control consisting of 20 pg of genomic DNA from each cell line used as input for scCUTseq. Neg cntr, negative control consisting of nuclease-free water. In all boxplots, each box spans from the 25th to the 75th percentile and whiskers extend from –1.5×IQR to +1.5×IQR from the closest quartile, where IQR is the inter-quartile range. Each dot corresponds to single well in the plate scheme shown below.

#### Supplementary Figure 2

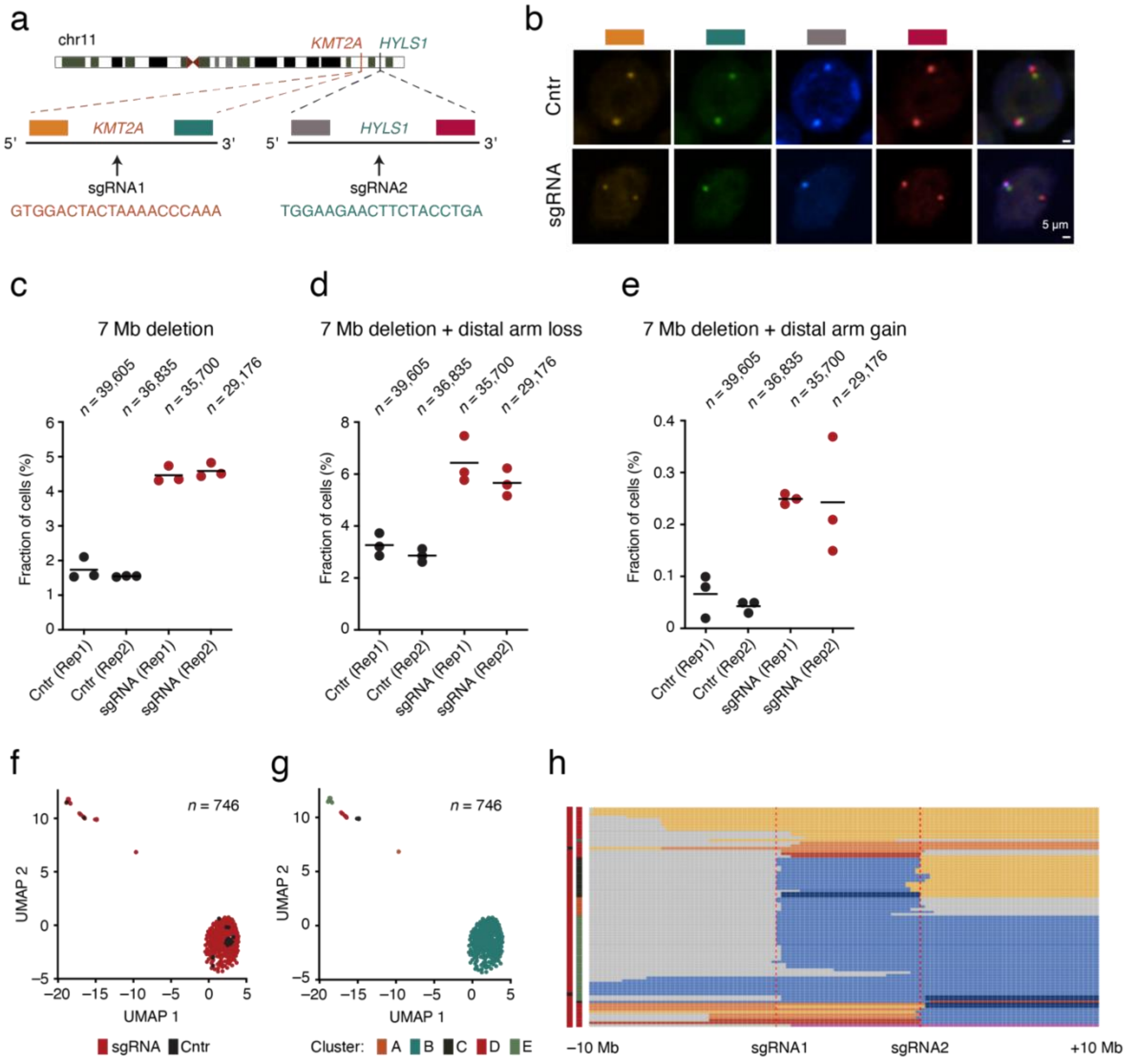


**Supplementary Fig. 2** scCUTseq sensitivity assessment. (**a**) Scheme of DNA fluorescence in situ hybridization (FISH) probes (colored rectangular bars) surrounding the *KMT2A* and *HYLS1* gene loci on chr11, used to detect a 7 Mb deletion induced by CRISPR-Cas9 using the two displayed small-guide RNAs (sgRNA). (**b**) Visualization of the FISH probes shown in (a) in human TK6 cells either transfected with non-targeting sgRNA (Cntr) or with both sgRNAs shown in (a) (sgRNA). The colored bars on the top correspond to the FISH probes shown in (a). In sgRNA treated cells in which the correct 7 Mb deletion event has occurred, the probes downstream of the *KMT2A* locus (green) and upstream of the *HYLS1* locus (grey) are detected only on one chr11 homologue, as expected. Scale bars, 5 µm. (**c**) Fraction of TK6 cells carrying the exact 7 Mb deletion on chr11. Each dot represents the fraction of cells containing the chromosomal rearrangement from a FISH technical replicate (one out of three wells of a 96-well plate). Horizontal black bars represent the mean. Rep, biological replicates. (**d**) Same as in (c), but for cells in which the 3′ portion of chr11 after the 7 Mb deletion was lost. (**e**) Same as in (c), but for cells in which the 3′ portion of chr11 after the 7 Mb deletion was amplified. (**f**) Dimensionality reduction by Uniform Manifold Approximation and Projection (UMAP) of copy number profiles of edited (sgRNA) and non-edited (Cntr) TK6 cells. Each dot represents a single cell. *n*, number of single cells. (**g**) Same as in (f), but with different clusters identified shown in different colors. (**h**) Single-cell copy number profiles (250 kb resolution) of TK6 cells harboring copy number alterations at the targeted location on chr11. The colors bars indicate edited (sgRNA) and non-edited (Cntrl) TK6 cells and different UMAP clusters, as in (f) and (g), respectively.

Supplementary Figure 3

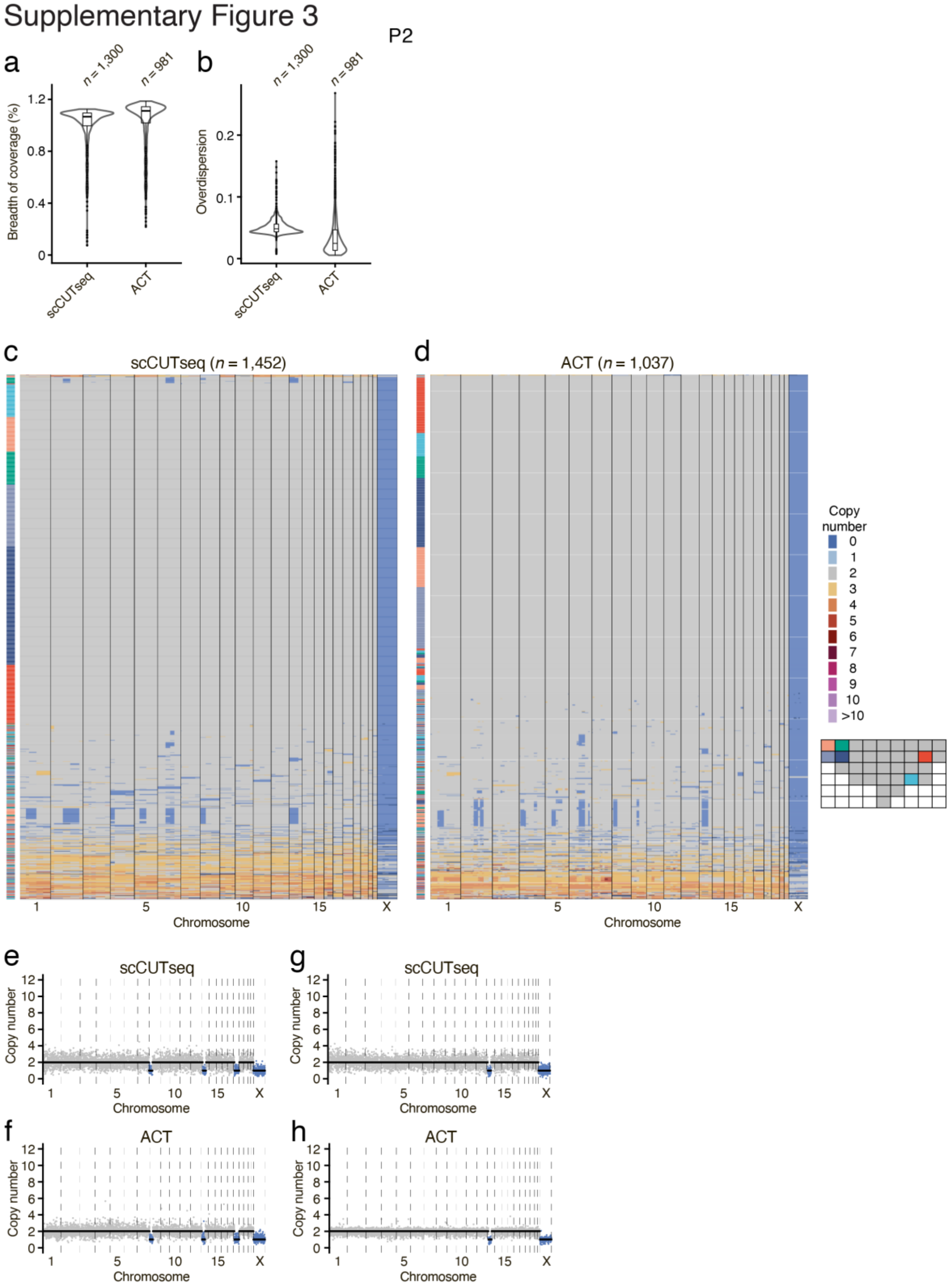


**Supplementary Fig. 3.** scCUTseq benchmarking. (**a**) Distributions of the breadth of genome coverage obtained with scCUTseq vs. Acoustic Cell Tagmentation (ACT) for the indicated number (*n*) of nuclei extracted from six different regions in prostate sample P2 (see scheme on the bottom right in (d). Violins extend from minimum to maximum, each box in the boxplot inside each violin spans from the 25th to the 75th percentile and whiskers extend from –1.5×IQR to +1.5×IQR from the closest quartile, where IQR is the inter-quartile range. Black dots, outliers. (**b**) Same as in (a) but for the overdispersion of binned read counts along the genome calculated as described in the Methods. (**c, d**) Single-cell copy number profiles (500 kb resolution) of the same cells profiled by scCUTseq (c) or ACT (d) analyzed in (a) and (b). The rectangular scheme on the bottom right in (d) shows the regions in prostate sample P3 from which the nuclei profiled by scCUTseq and ACT were obtained. See **Fig. 1c** for the corresponding histopathologic annotation. (**e-h**) Examples of copy number profiles (500 kb resolution) of pseudo-diploid cells displaying the same deletion patterns detected by both scCUTseq and ACT (compare (e) with (f) and (g) with (h)). Each plot corresponds to one cell.

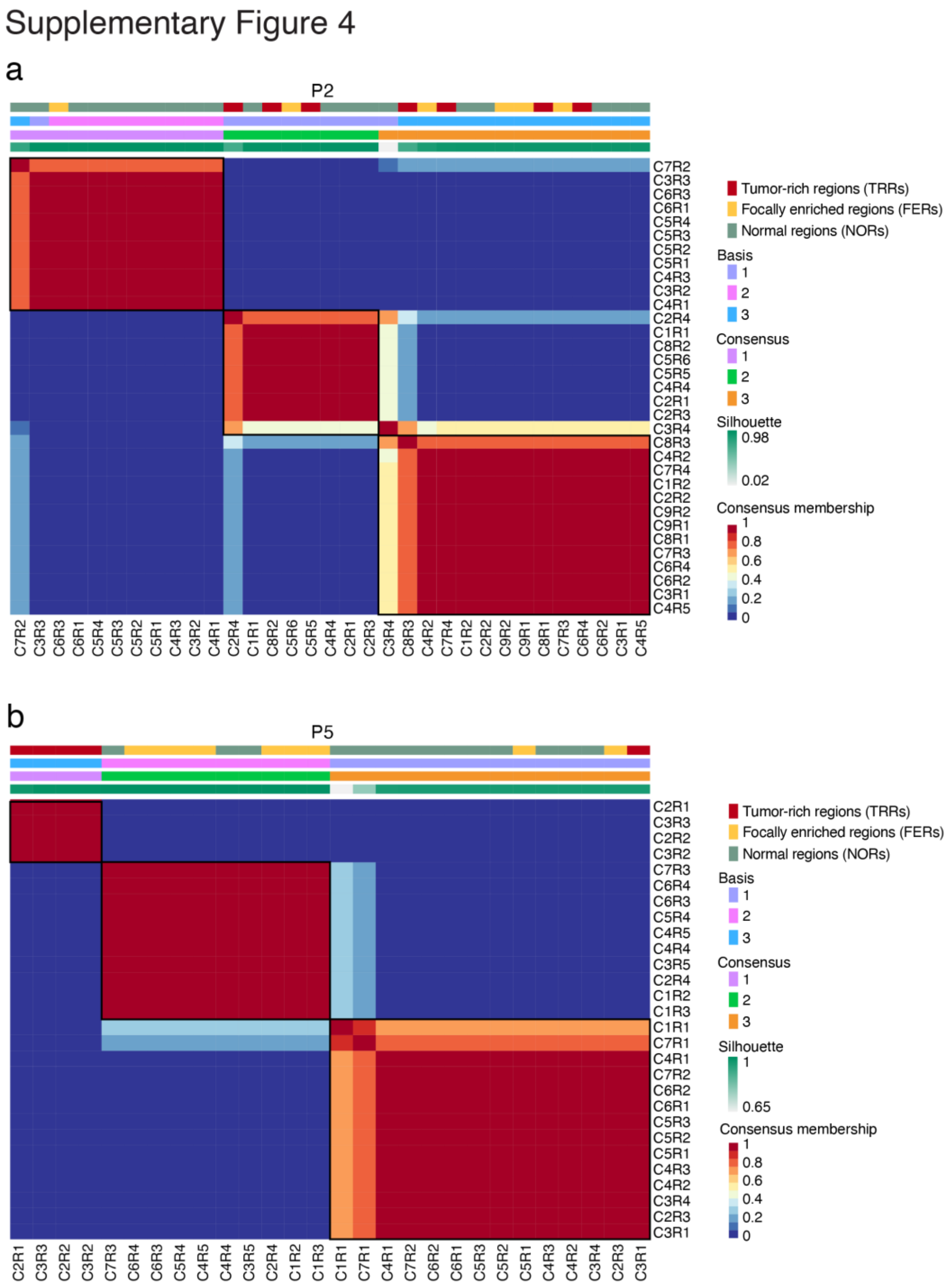


**Supplementary Fig. 4.** Annotation by RNA-seq of tissue regions in prostate samples P2 and P5. See **Supplementary Methods** for how RNA-seq data clustering was performed. (**a**) Quality of clustering of the regions in P2 for *n*=3 clusters corresponding to the number of different region types defined based on histophatology. The position of each region in the corresponding tissue map in **Fig. 1c** is indicated (C, column; R, row). (**b**) Same as in (a) but for sample P5. The position of each region in the corresponding tissue map in **Fig. 1d** is indicated.

Supplementary Figure 5

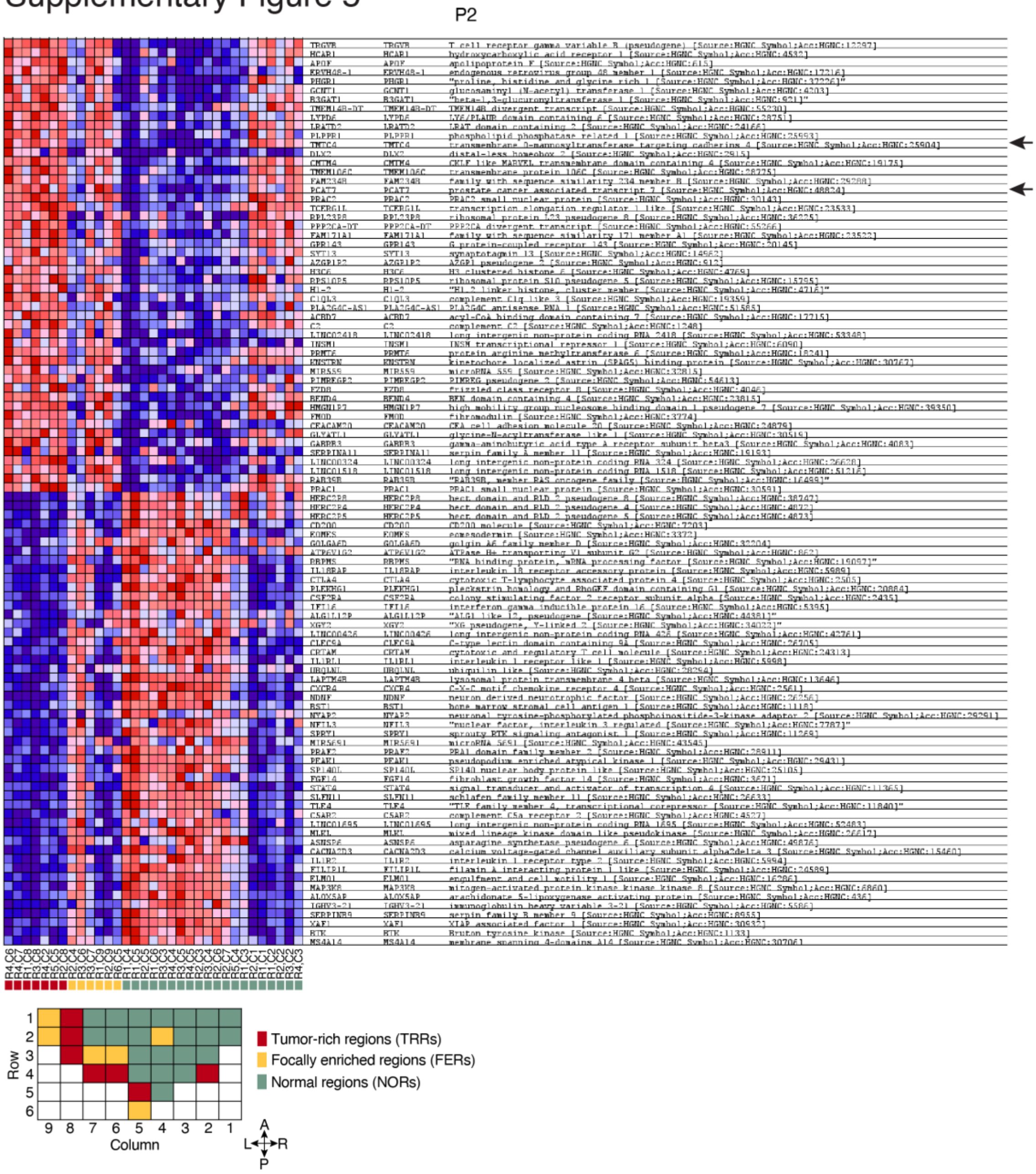


**Supplementary Fig. 5.** Gene set enrichment analysis (GSEA) comparing tumor-rich regions (TRRs) versus focally enriched regions (FERs) and normal regions (NORs) in prostate sample P2. The top 50 up- and down-regulated genes are shown. The arrows on the right indicate two prostate cancer biomarker genes (*PCA3* and *TMTC4*) that are upregulated in TRRs. The tissue map on the bottom left is the same as in **Fig. 1c**.

### Supplementary Figure 6

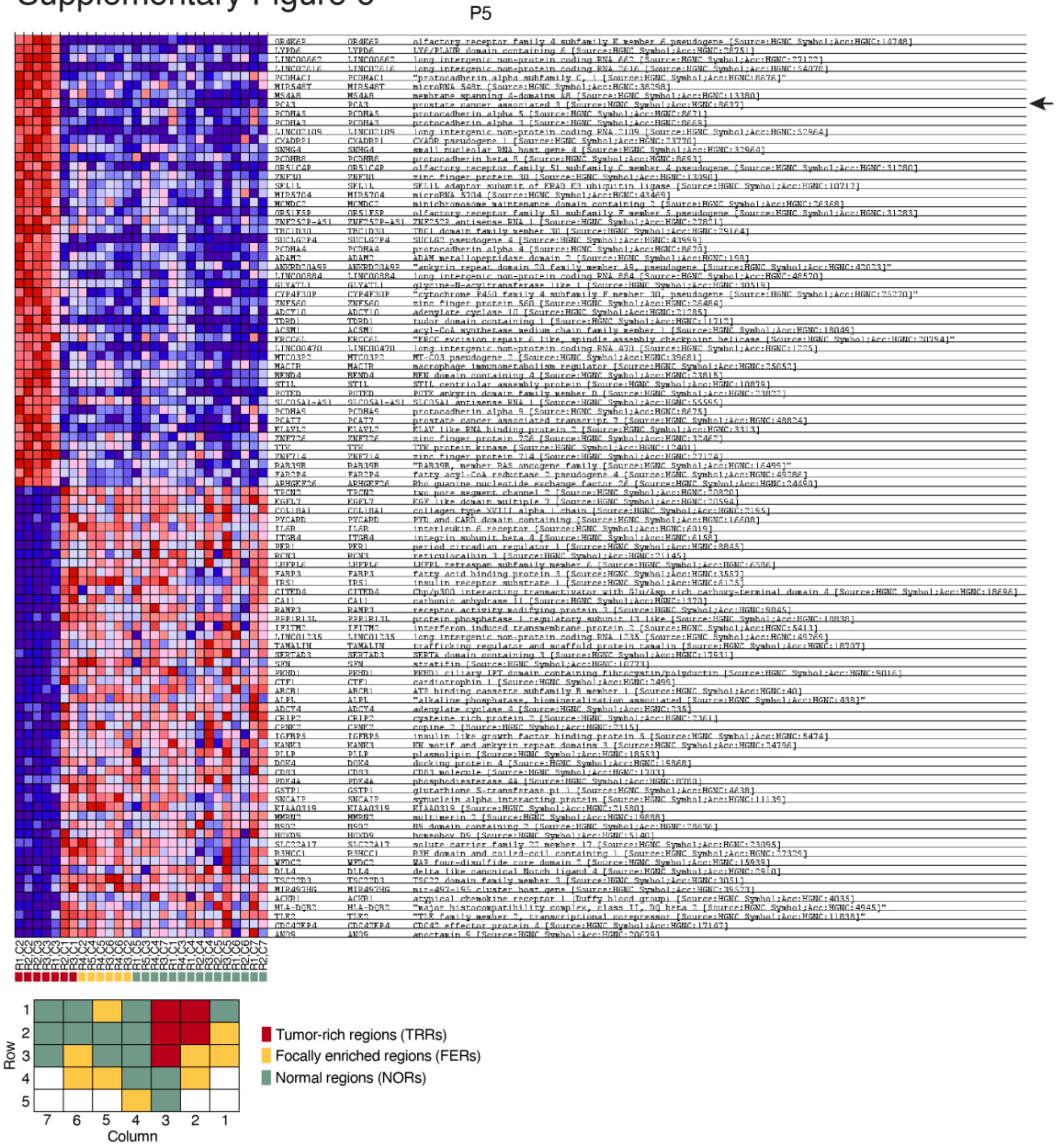


**Supplementary Fig. 6.** Gene set enrichment analysis (GSEA) comparing tumor-rich regions (TRRs) versus focally enriched regions (FERs) and normal regions (NORs) in prostate sample P5. The top 50 up- and down-regulated genes are shown. The arrow on the right indicates a prostate cancer biomarker gene (*PCA3*) that is upregulated in TRRs. The tissue map on the bottom left is the same as in **Fig. 1c**.

Supplementary Figure 7

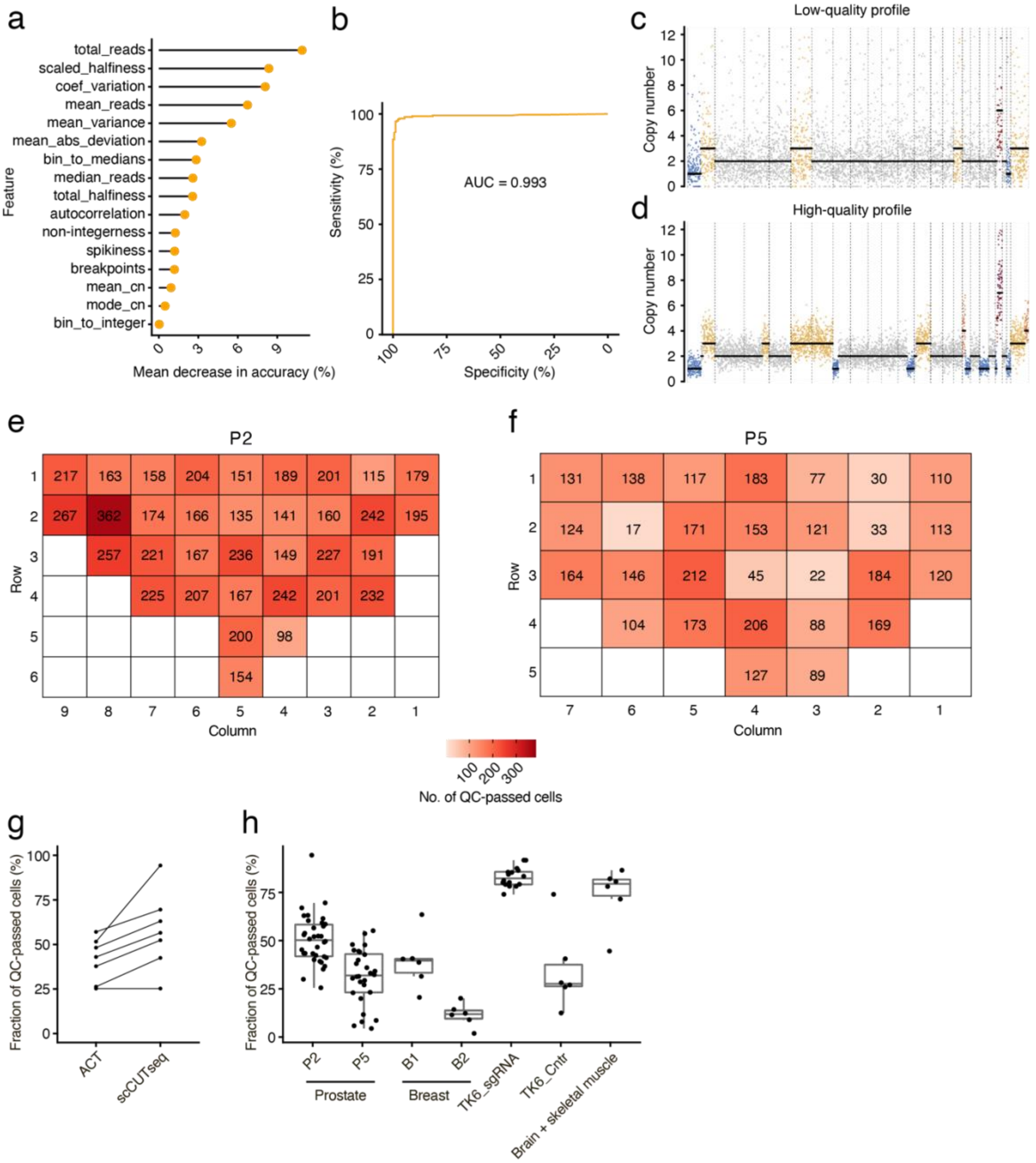


**Supplementary Fig. 7.** Random forest classifier of single-cell copy number profiles. (**a**) Ranked importance of the features used by the random forest classifier trained on scCUTseq data to distinguish between high- and low-quality copy number profiles. See **Supplementary Table 3** for a description of each feature. (**b**) Receiver operating characteristic curve analysis of the random forest classifier performance. AUC, area under the curve. (**c, d**) Examples of low-quality (c) and high-quality (d) quality scCUTseq copy number profiles (500 kb resolution). Low-quality profiles are discarded by the random forest classifier, whereas high-quality profiles are retained for further analysis. Each dot represents a 500 kilobases (kb) genomic bin. Black, blue, yellow, and red dots indicate, respectively, copy number levels determined by circular binary segmentation, chromosomal deletions, gains, and amplifications. (**e, f**) Heatmaps displaying the number of cells with high-quality scCUTseq copy number profiles retained by the random forest classifier, for each region in prostate samples P2 (e) and P5 (f) profiled by scCUTseq. (**g**) Comparison of the fraction of cells with high-quality copy number profiles that passed quality control (QC) by the random forest classifier, for six regions profiled by both ACT and scCUTseq in prostate sample P2 (see **Supplementary Fig. 3d** for a map of the regions). (**h**) Distributions of the fraction of cells with high-quality copy number profiles obtained in different scCUTseq experiments performed on different cell/tissue types. Copy number profiles of brain and breast samples are shown in **Supplementary Fig. 8 and 12**, respectively. TK6_sgRNA and TK6_Cntr cells are described in **Supplementary Fig. 2**. In all boxplots, each box spans from the $25^{th}$ to the $75^{th}$ percentile and whiskers extend from –1.5×IQR to +1.5×IQR from the closest quartile, where IQR is the inter-quartile range. Each dot in the boxplots represents one library (see **Supplementary Table 9** for a list of all libraries sequenced in this study).

Supplementary Figure 8

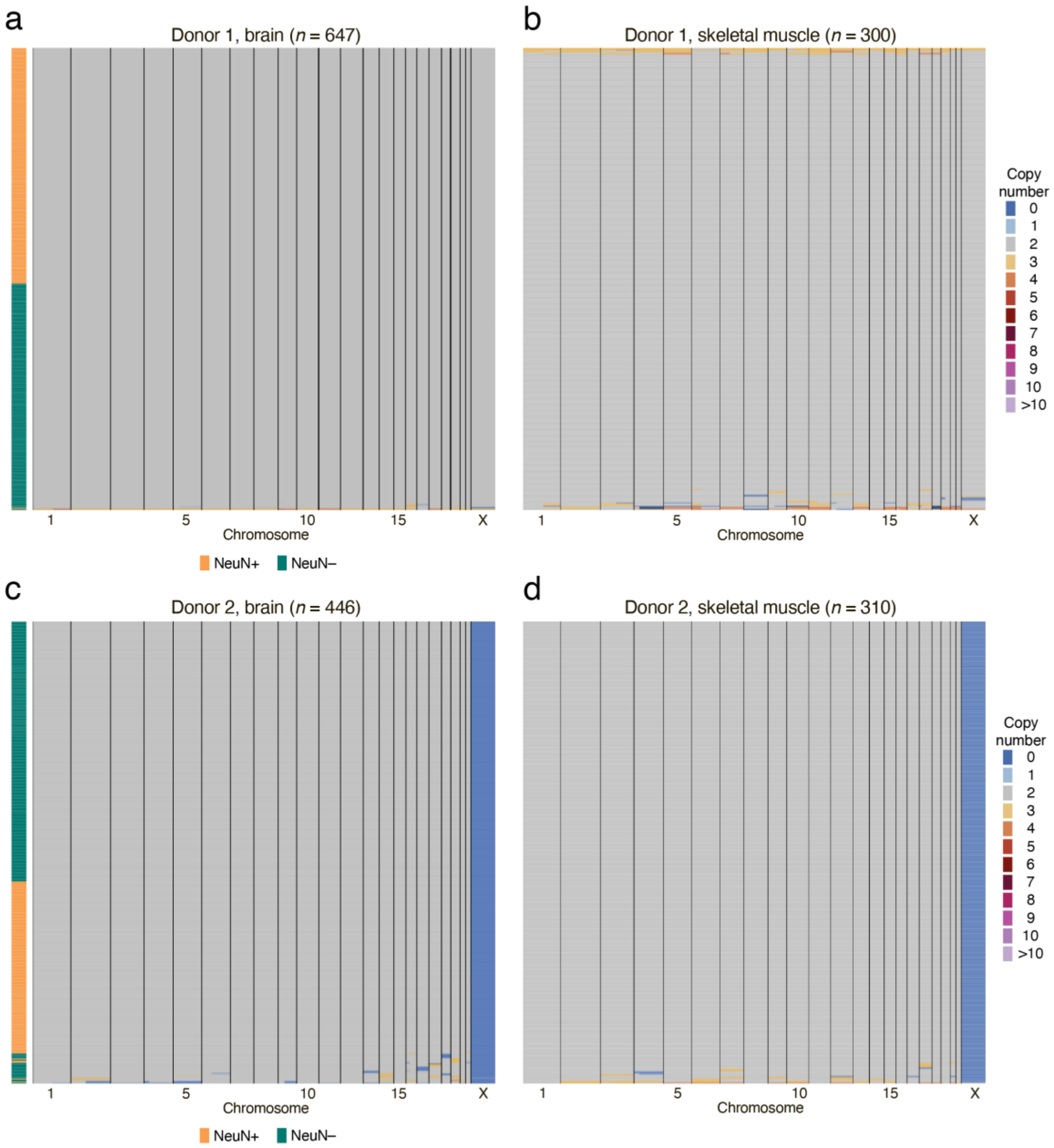


**Supplementary Fig. 8.** Absence of gross copy number artefacts in scCUTseq. (**a, b**) Single-cell copy number profiles (500 kb resolution) obtained by applying scCUTseq to nuclei extracted from pre-frontal cortex (brain) and skeletal muscle from one donor in the Donatum autopsy program (see **Methods and Supplementary Table 4**). Brain nuclei were sorted in neuronal (NeuN+) and non-neuronal (NeuN–) populations by FACS. *n*, number of single cells displayed. As it can be seen, the vast majority of nuclei sequenced shows a diploid copy number state, with very few cells displaying copy number alterations, indicating that the whole-genome amplification step in scCUTseq does not cause overt copy number artefacts. (**c, d**) Same as in (a, b) but for a different donor.

Supplementary Figure 9

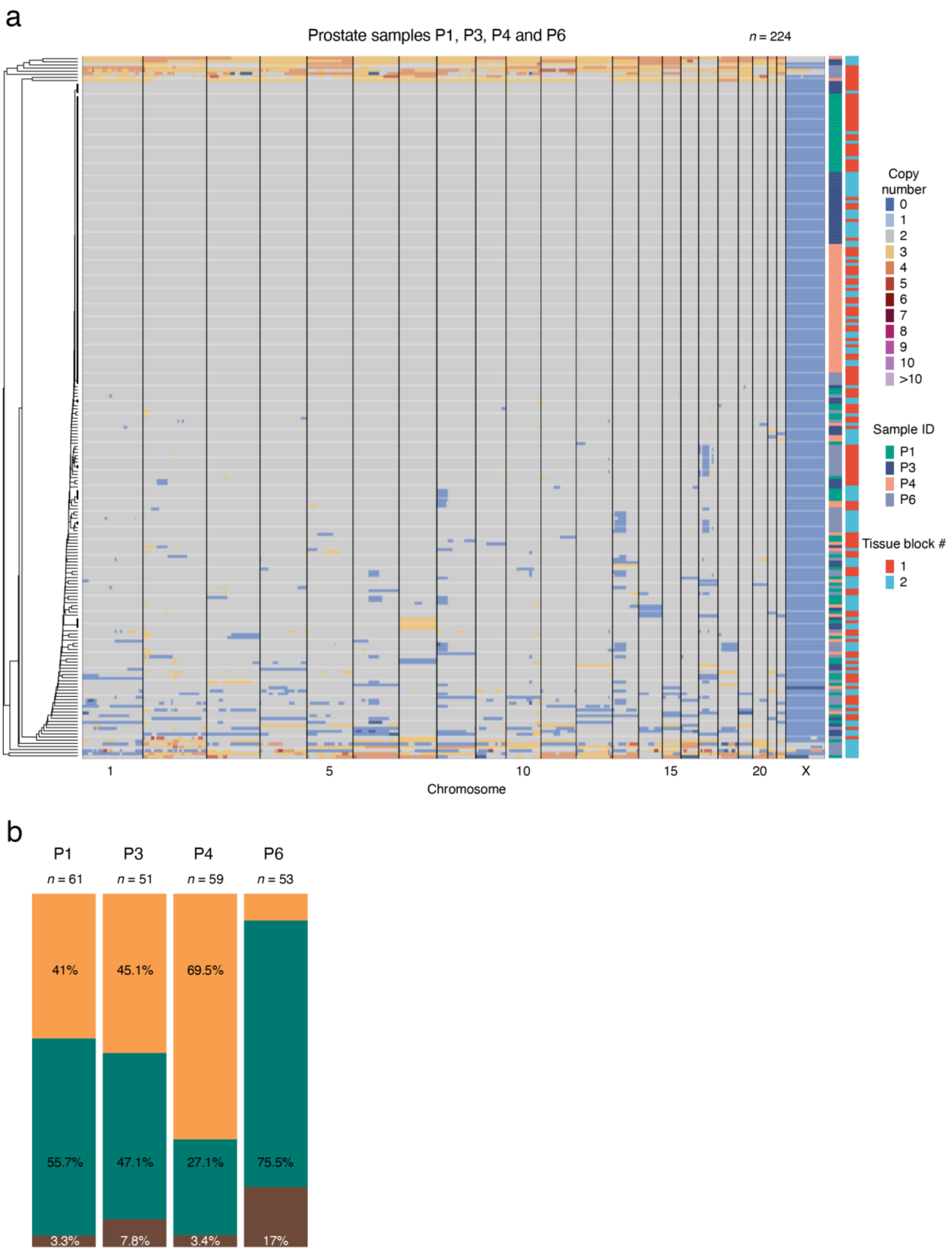


**Supplementary Fig. 9.** Single-cell copy number profiles (500 kb resolution) for the other four prostatectomy samples (P1, P3, P4, P6) that were profiled by scCUTseq. Two regions (tissue blocks) were profiled in each sample. (**a**) Hierarchically clustered scCUTseq profiles from all the cells sequenced in the four samples. (**b**) Fraction of diploid, ‘pseudo-diploid and ‘monster’ cells in the four samples shown in (a). *n*, number of single cells with high-quality copy number profiles analyzed.

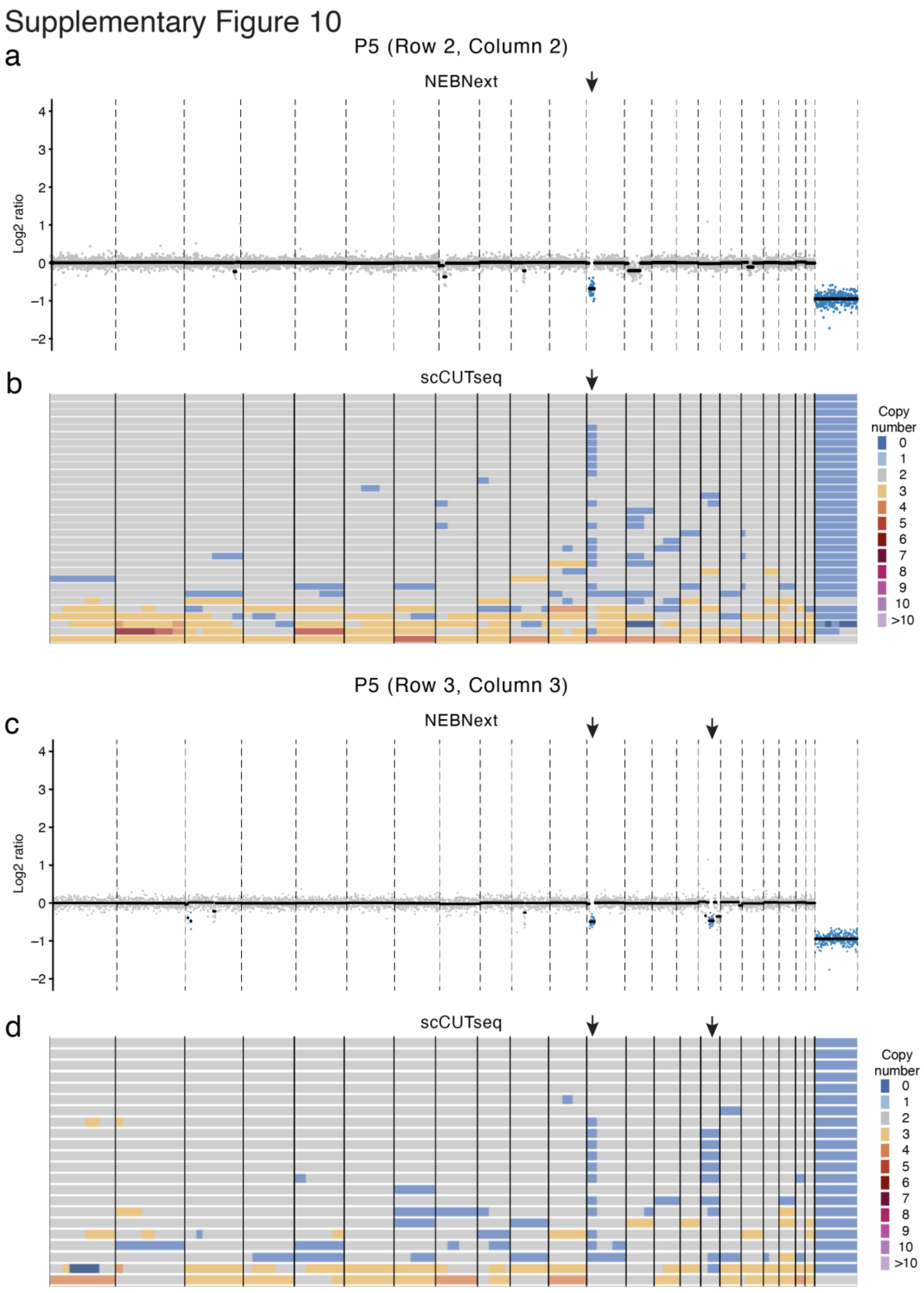


**Supplementary Fig. 10.** Bulk sequencing only detects subclonal copy number changes present in many cells. (**a, b**) Bulk copy number profile (500 kb resolution) from gDNA extracted from a single region in prostate sample P2 and used for library preparation using a commercial kit (NEBNext) (a) and corresponding single-cell copy number profiles (500 kb resolution)

obtained by applying scCUTseq to nuclei extracted from the same region (b). The vertical arrows pinpoint a subclonal deletion that is present in many cells and is therefore also detected by bulk DNA-seq. See **Fig. 1c** for the location of the region in sample P2. (**c, d**) Same as in (a, b) but for a different region in sample P2.

Supplementary Figure 11

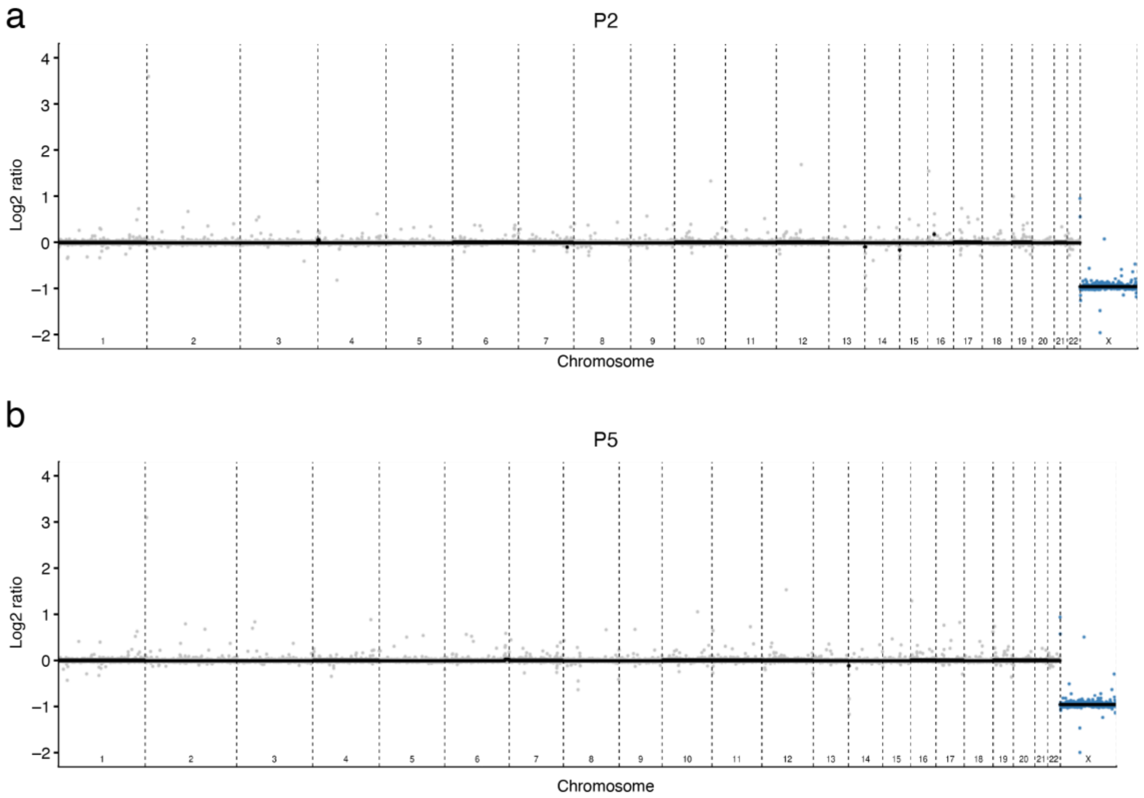


**Supplementary Fig. 11.** Absence of germline copy number variants in the donors of prostate samples P2 and P5 profiled by scCUTseq. Libraries for whole-genome sequencing (WGS) were prepared from peripheral blood gDNA using a commercial kit (NEBNext), as described in the **Supplementary Methods**. Each gray dot corresponds to a 250 kb genomic bin.

Supplementary Figure 12

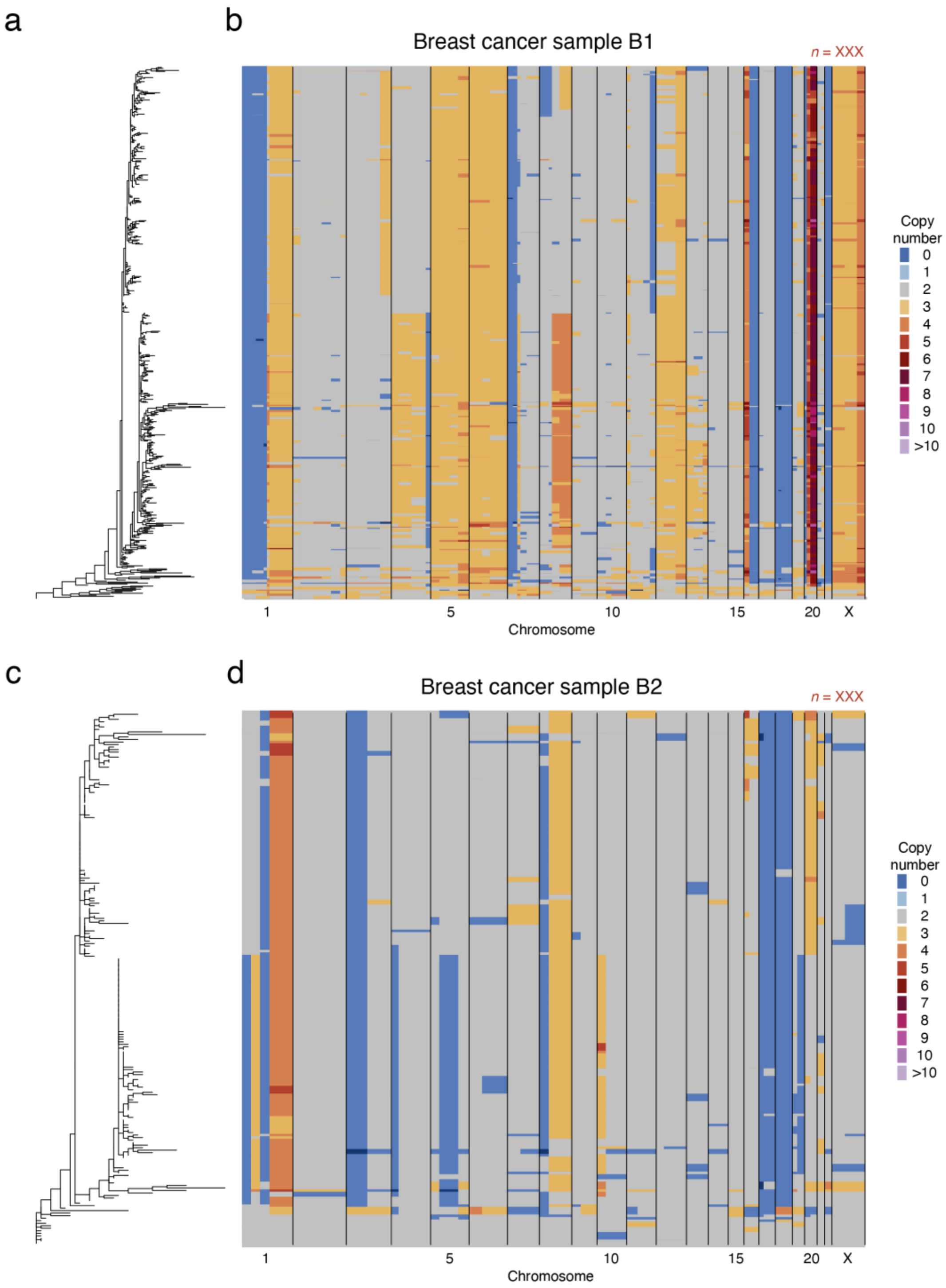


**Supplementary Fig. 12.** (**a**) Phylogenetic Newick tree of single cells extracted from a breast cancer surgical biopsy and profiled by scCUTseq (see **Methods**). The tree was generated by *MEDICC2*[19] (see **Methods**). Each leaf in the tree corresponds to one cell. (**b**) Copy number profiles (500 kb resolution) of the cells shown in (b). (**c, d**) Same as in (a, b) but for another breast cancer specimen.

Supplementary Figure 13

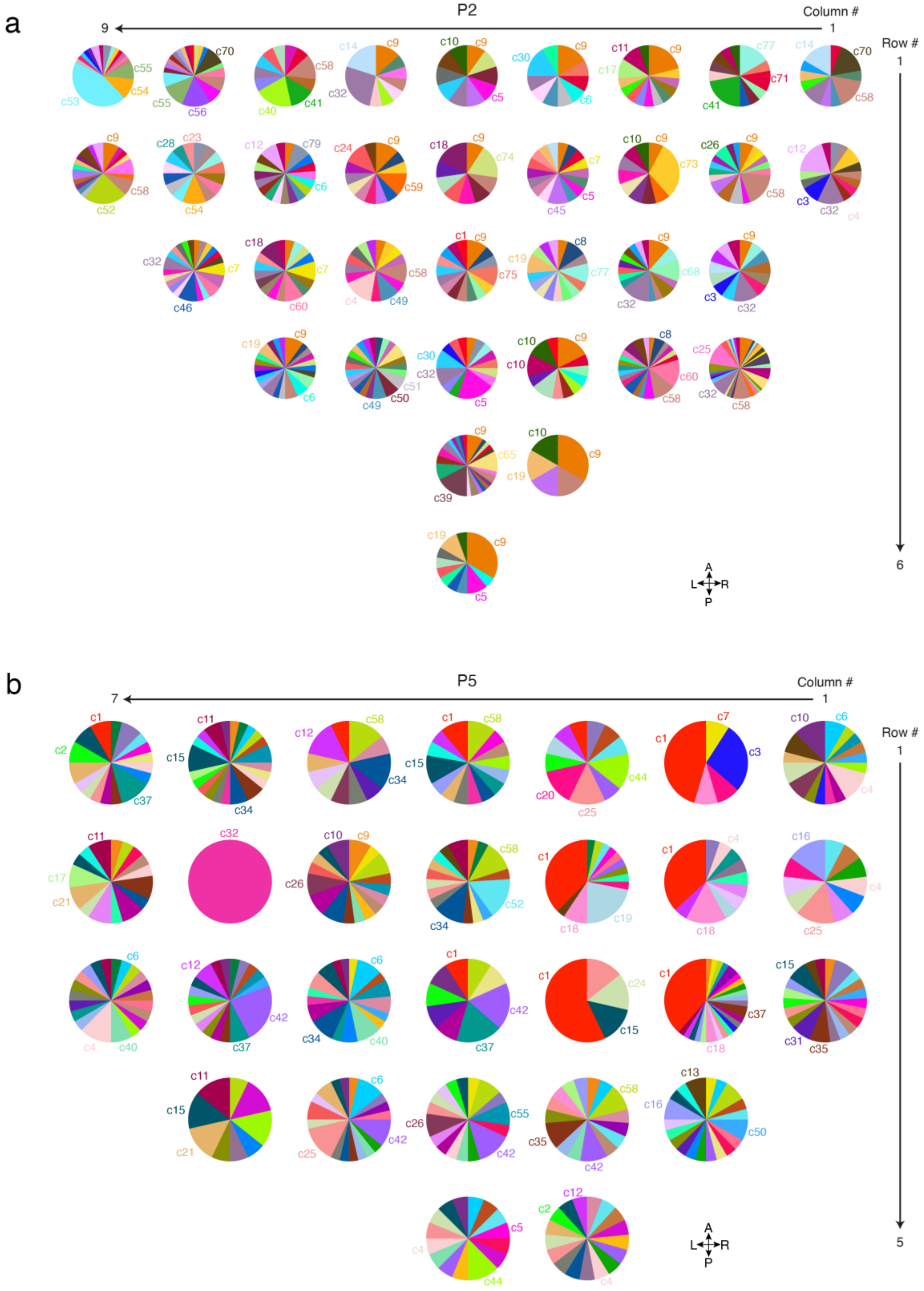


**Supplementary Fig. 13.** Proportion of pseudo-diploid subclones in each of the regions profiled by scCUTseq in prostate samples P2 (**a**) and P5 (**b**). Each pie chart corresponds to one tissue cube (region). The three most frequent subclones (c) in each tissue block are indicated. The anatomical orientation of the tissues is shown by the four arrows on the bottom right. A, anterior. P, posterior. L, left. R, right. Rows and columns numbers are the same as in **Fig. 1c, d**.

Supplementary Figure 14

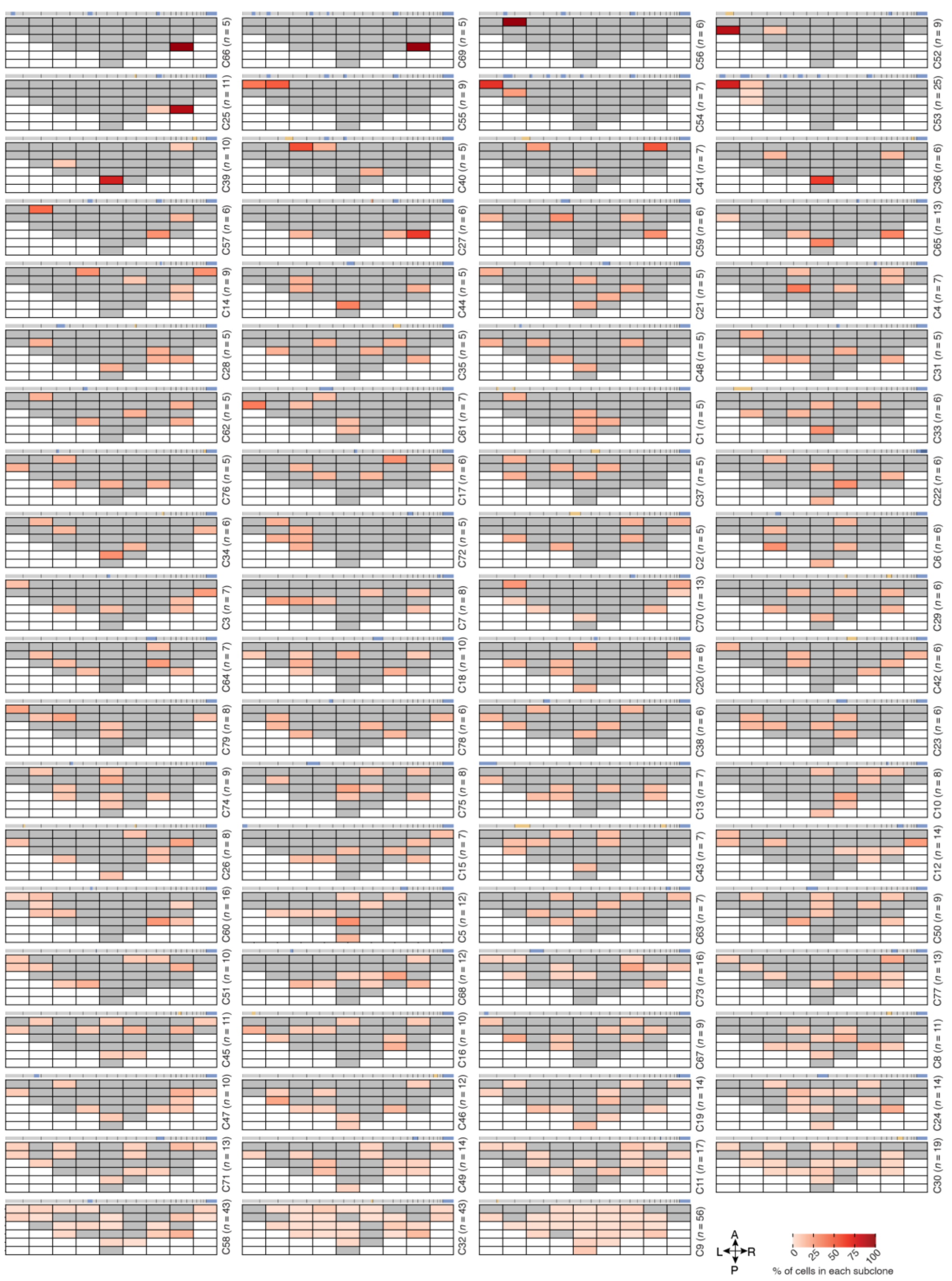


**Supplementary Fig. 14.** Spatial distribution of the pseudo-diploid subclones (C) identified in prostate sample P2. *n*, number of pseudo-diploid cells in each subclone. Each grid represents the tissue distribution of the corresponding subclone (indicated on the right side) and the regions in which the subclone was found are color-coded based on the percentage of cells belonging to that subclone. Grey cells indicate tissue regions in which the corresponding subclone was not detected. White cells indicate absence of prostate tissue. The bar above each grid shows the genome-wide median copy number profile of the corresponding subclone. Blue bars indicate deletions, orange bars amplifications. Vertical black bars mark the boundaries between consecutive chromosomes. The anatomical orientation of the grids is shown by the four arrows on the bottom right. A, anterior. P, posterior. L, left. R, right.

Supplementary Figure 15

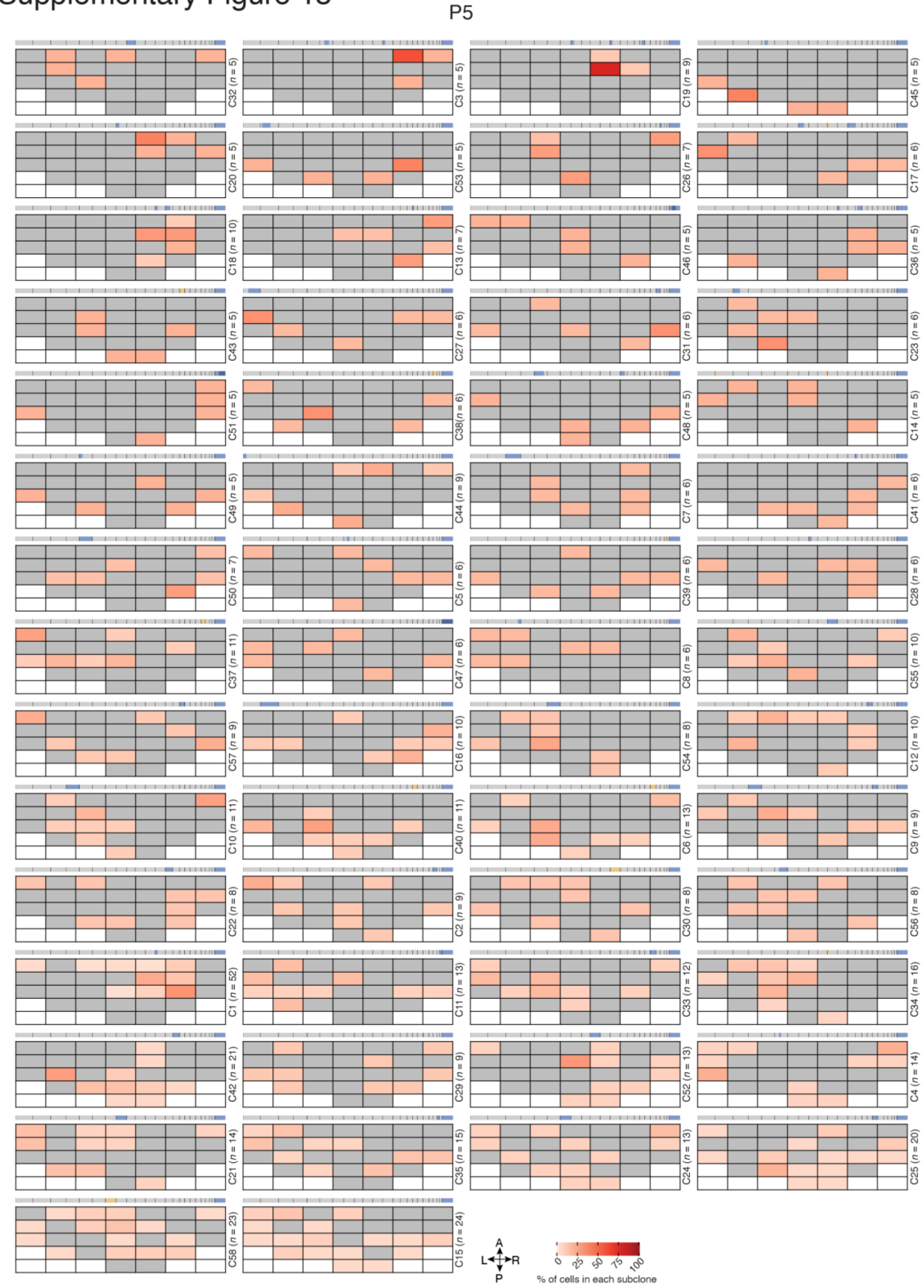


**Supplementary Fig. 15.** Spatial distribution of the pseudo-diploid subclones (C) identified in prostate sample P5. *n*, number of pseudo-diploid cells in each subclone. Each grid represents the tissue distribution of the corresponding subclone (indicated on the right side) and the regions in which the subclone was found are color-coded based on the percentage of cells belonging to that subclone. Grey cells indicate tissue regions in which the corresponding subclone was not detected. White cells indicate absence of prostate tissue. The bar above each grid shows the genome-wide median copy number profile of the corresponding subclone. Blue bars indicate deletions, orange bars amplifications. Vertical black bars mark the boundaries between consecutive chromosomes. The anatomical orientation of the grids is shown by the four arrows on the bottom right. A, anterior. P, posterior. L, left. R, right.

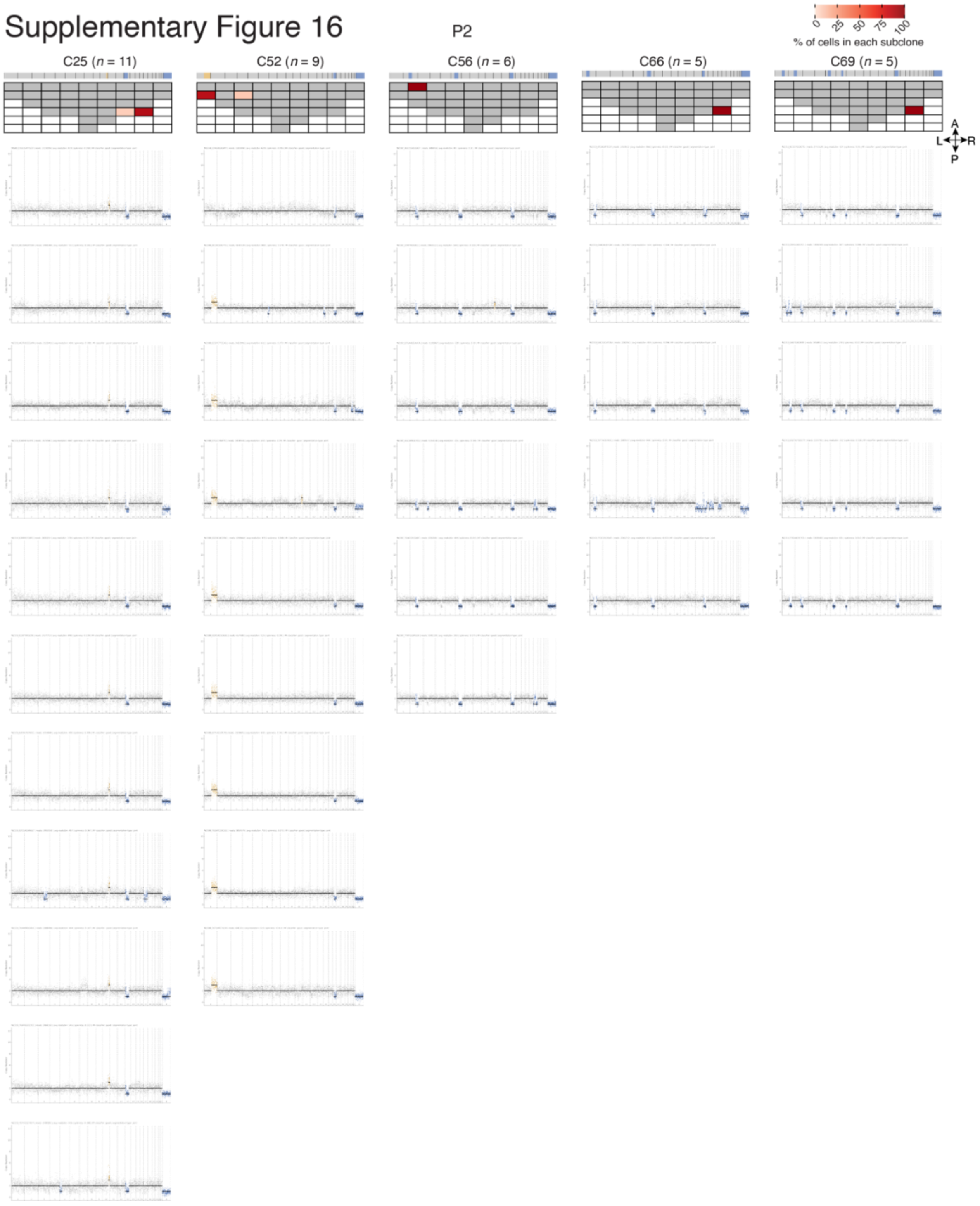


**Supplementary Fig. 16.** Individual copy number profiles (500 kb resolution) of the pseudo-diploid cells in each of the five most localized subclones (C) identified in prostate sample P2. *n*, number of pseudo-diploid cells in each subclone. In each copy number plot, dots represent 500 kb genomic bins. Orange and blue dots indicate, respectively, amplifications and deletions. Grey dots indicate bins belonging to normal (diploid) genomic segments. The grids on the top represent the tissue distribution of the corresponding subclone. Grey cells indicate tissue regions in which the corresponding subclone was not detected. White cells indicate absence of prostate tissue. The bar above each grid shows the genome-wide median copy number profile of the corresponding subclone. The anatomical orientation of the grids is shown by the four arrows on the right. A, anterior. P, posterior. L, left. R, right.

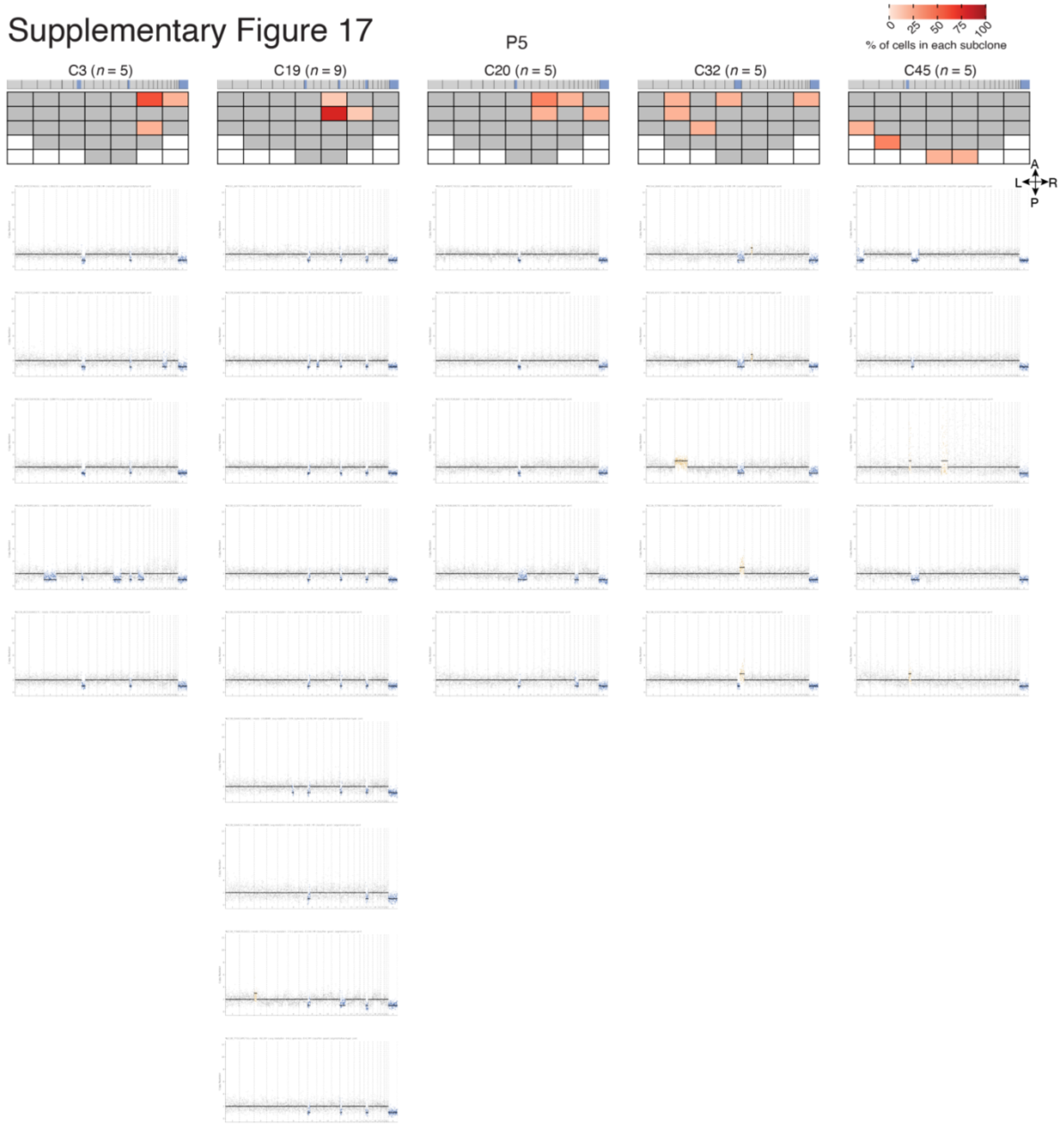


**Supplementary Fig. 17.** Individual copy number profiles (500 kb resolution) of the pseudo-diploid cells in each of the five most localized subclones (C) identified in prostate sample P5. *n*, number of pseudo-diploid cells in each subclone. In each copy number plot, grey dots represent 500 kb genomic bins. Orange and blue dots indicate, respectively, amplifications and deletions. Grey dots indicate bins belonging to normal (diploid) genomic segments. The grids on the top represent the tissue distribution of the corresponding subclone. Grey cells indicate tissue regions in which the corresponding subclone was not detected. White cells indicate absence of prostate tissue. The bar above each grid shows the genome-wide median copy number profile of the corresponding subclone. The anatomical orientation of the grids is shown by the four arrows on the right. A, anterior. P, posterior. L, left. R, right.

Supplementary Figure 18

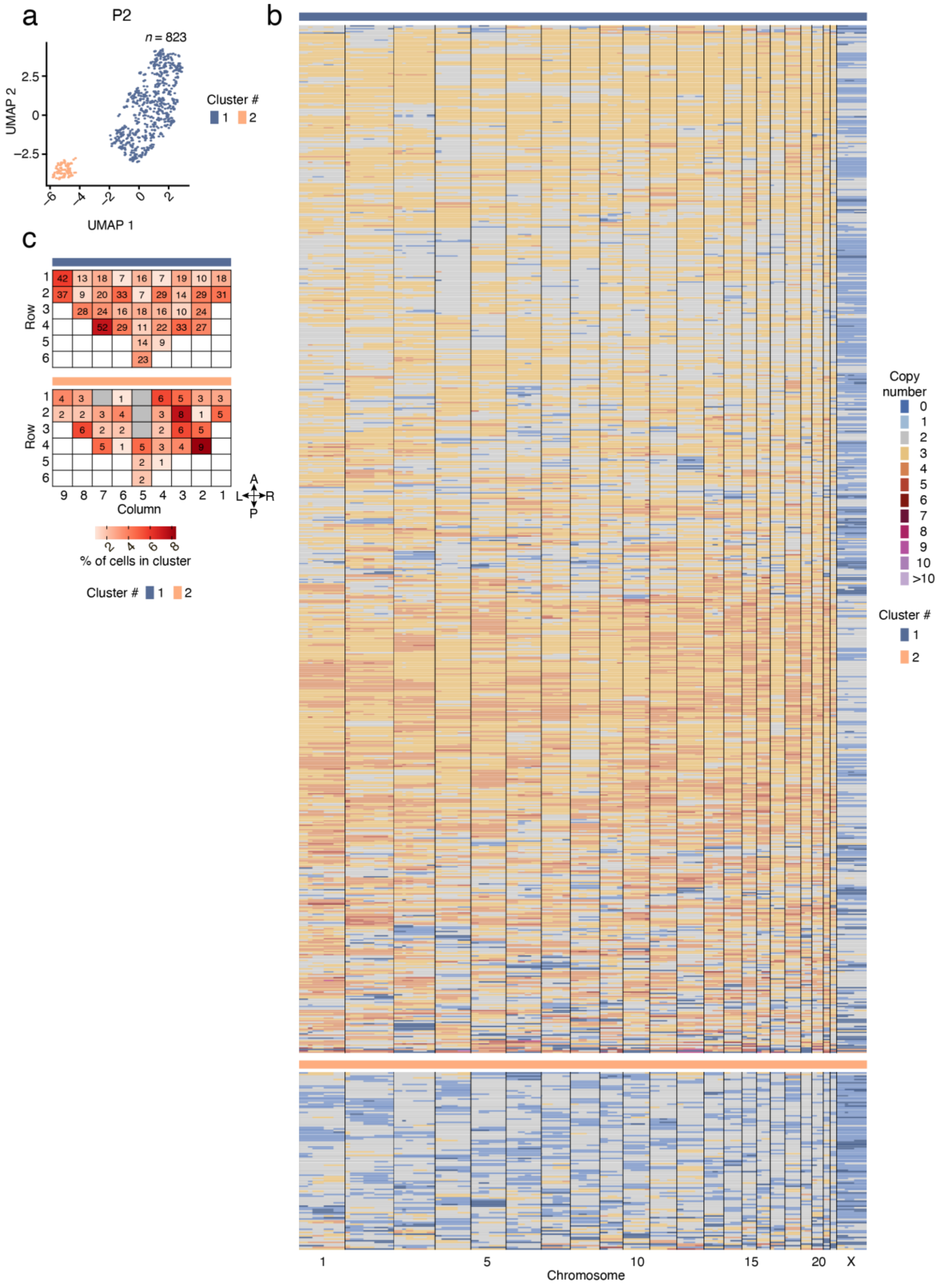


**Supplementary Fig. 18.** Characteristics and spatial distribution of monster cells in prostate sample P2. (**a**) Uniform Manifold Approximation and Projection (UMAP) dimensionality reduction of monster cell SCNA profiles. *n*, number of cells. Dots of the same color belong to the same cluster. Each dot represents a single cell. (**b**) Single-cell copy number profiles of monster cells in each of the two UMAP clusters shown in (a). (**c**) Maps showing the tissue distribution of monster cells for each of the two UMAP clusters in (a). In each grid, each cell represents a tissue region from which nuclei were isolated and profiled by scCUTseq. The numbers in each cell of the grids indicate the number of monster cells assigned to the indicated cluster in that region. Grey cells mark tissue regions in which no monster cell was detected. White cells indicate absence of prostate tissue. The anatomical orientation of the maps is shown by the four arrows on the bottom right. A, anterior. P, posterior. L, left. R, right.

#### Supplementary Figure 19

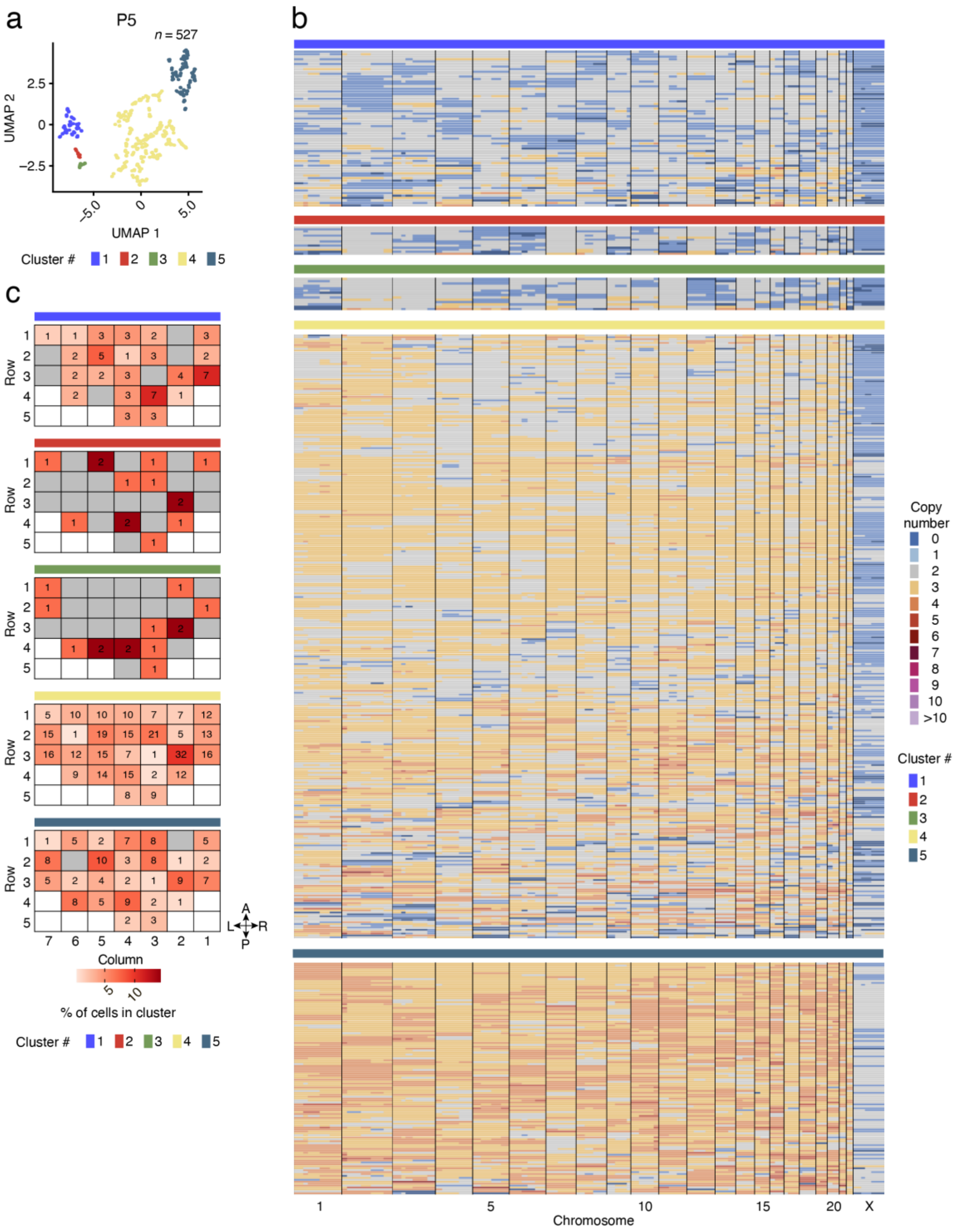


**Supplementary Fig. 19.** Characteristics and spatial distribution of monster cells in prostate sample P5. (**a**) Uniform Manifold Approximation and Projection (UMAP) dimensionality reduction of monster cell SCNA profiles. *n*, number of cells. Dots of the same color belong to the same cluster. Each dot represents a single cell. (**b**) Single-cell copy number profiles of monster cells in each of the two UMAP clusters shown in (a). (**c**) Maps showing the tissue distribution of monster cells for each of the two UMAP clusters in (a). In each grid, each cell represents a tissue region from which nuclei were isolated and profiled by scCUTseq. The numbers in each cell of the grids indicate the number of monster cells assigned to the indicated cluster in that region. Grey cells mark tissue regions in which no monster cell was detected. White cells indicate absence of prostate tissue. The anatomical orientation of the maps is shown by the four arrows on the bottom right. A, anterior. P, posterior. L, left. R, right.

Supplementary Figure 20

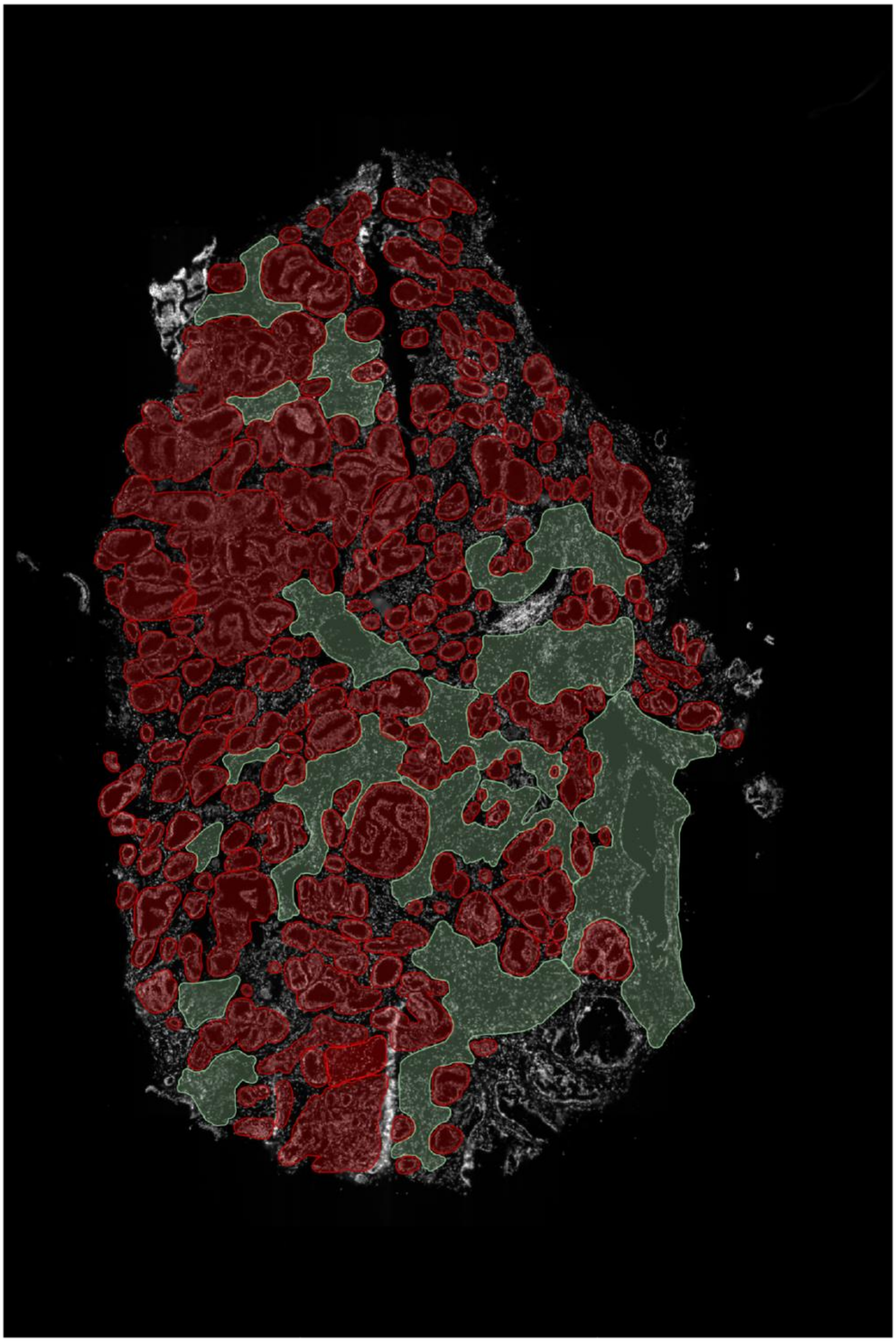

**Supplementary Fig. 20.** QuPath annotation of a tissue section adjacent to the one profiled by DNA FISH in prostate sample P6 (see **Fig. 3**). Red, neoplastic prostate glands. Green, stroma.

#### Supplementary Figure 21

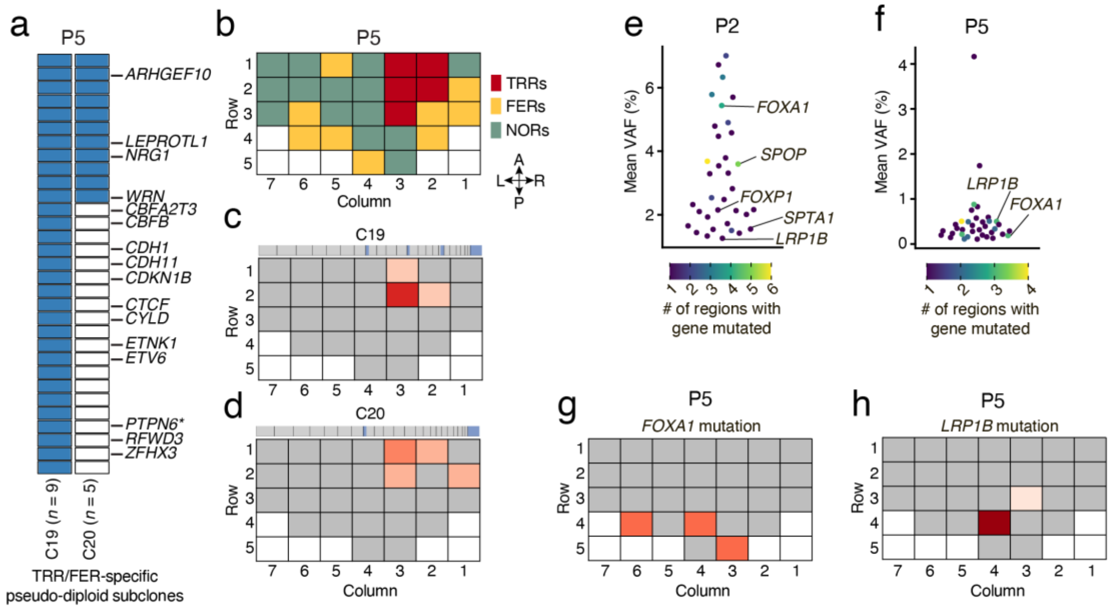


**Supplementary Fig. 21.** Loss of tumor-suppressor genes and mutations in prostate cancer associated genes in prostate sample P5. (**a**) OncoPrint plot showing genes classified as tumor-suppressor genes (TSGs) in the Catalogue of Somatic Mutations in Cancer (COSMIC) that were deleted (blue rectangles) in two pseudo-diploid subclones (C) localized exclusively in tumor-rich regions (TRRs) or focally enriched regions (FERs) in prostate sample P5. *n*, number of pseudo-diploid cells in each subclone. Asterisks indicate genes annotated in COSMIC both as TSGs and as oncogenes (context-dependent TSGs). (**b**) Same as in **Fig. 1d**. (**c, d**) Spatial distribution of the two pseudo-diploid subclones (C) shown in (a). The median copy number profile of each subclone is shown on top of the corresponding tissue map. (**e, f**) Mean variant allelic frequency (VAF) of cancer-associated genes included in the TSO500 panel (Illumina) across all regions in which the genes were found mutated (non-synonymous SNVs) in prostate samples P2 and P5, respectively. Each dot corresponds to one gene. Genes previously associated with prostate cancer are labeled. (**g, h**) Schematic maps showing where the indicated genes were found mutated in sample P2. Grey cells indicate tissue regions in which the indicated gene was not altered. White cells indicate absence of tissue. The anatomical orientation of all the maps in (c, d) and (g, h) is the same as in (b).

#### 2. Supplementary Methods

**RNA-seq**

We prepared RNA-seq libraries from the total RNA extracted from each region in the P3 and P6 prostate samples (see **Supplementary Table 1**) using the TruSeq Stranded Total RNA Library Prep kit (Illumina, cat. no. 20020597) with an RNA input ranging from 800 ng to 1 μg of total RNA with RIN > 6. After removal of rRNA, we fragmented the RNA for 5 min, followed by cDNA synthesis, end-repair, 3'-adenylation, and adapter ligation following the manufacturer's instructions. Lastly, we sequenced pooled libraries on Illumina NovaSeq 6000 with a loading concentration equal to 1.2 nM and paired-end 100 bp sequencing mode.

To analyze the data, we aligned paired-end reads to the GRCh38 genome assembly using the *HISAT2* aligner[1]. We filtered the resulting bam files for ribosomal RNA sequences using the *RSeQC* module *split_bam.by*. We estimated gene expression abundance using the *FeatureCounts* function in the *Rsubread* (version 2.14.2)[2] R package and GENCODE (v40) genomic features annotation. We then subjected gene expression counts to filtering of lowly expressed features, trimmed-mean of M-values (TMM) normalization, and counts per million (CPM) calculation using the *edgeR* (version 3.42.4)[3] R package. For class discovery, we used a consensus based non-negative matrix factorization (NMF) approach implemented in the *NMF* (version 0.26)[4] R package. We carried out unsupervised count matrix decomposition on the top 25% most variable features for increasing number of factorization ranks ranging from 2 to 6 selecting 200 iterations. Lastly, we used the Gene Set Enrichment Analysis (GSEA) software (version 4.2.3)[5] and human *MSigDB c2.all.v2023.1.Hs.symbols.gmt* gene set to extract biomarkers for each supervised cluster (i.e., TRRs, FERs, and NORs).

**Standard MALBAC**

We washed the harvested cells in 1× PBS/5 mM EDTA at room temperature and resuspended them in the same buffer at a density of $10^6$/mL. We then placed the cells either on ice for immediate sorting (live cells) or fixed them to store them for longer periods before sorting. For fixation, we added an equal volume of 1× PBS/5 mM EDTA/8% methanol-free paraformaldehyde (PFA, Thermo Fisher Scientific, cat. no. 28908) to the cell suspension and pipetted the solution up and down ten times. After 10 min incubation in darkness, we added 2.5 M glycine to each tube to reach a final concentration of 125 mM to quench any residual unreacted PFA. We then washed the cells in 1× PBS/5 mM EDTA at room temperature, resuspended them at a density of $10^6$/mL in the same buffer, and stored them at +4 °C in 1× PBS/5 mM EDTA/0.05% $NaN_3$. We have successfully used fixed cells prepared in this manner, that were kept at +4 °C several weeks up to three months. Before sorting, we transferred the cells into FACS-compatible tubes, stained them with 2.46 ng/mL Hoechst 33342 (Thermo Fisher Scientific, cat. no. 62249) and incubated for 40 min at 37 °C in darkness, while rotating. We sorted fixed and live single cells in a 96-well plate pre-filled with MALBAC lysis reaction mix (5 µL of lysis buffer + 0.5 µL of lysis enzyme) using a BD FACSJazz Cell Sorter (BD Biosciences) based on forward and side scatter properties. We centrifuged the plate at 600×g for 3 min. In parallel we used one well as negative control by adding 1 µL of Nuclease-Free Water (Thermo Fisher Scientific, cat. no. 4387936) into the lysis mix and another well as positive control by adding 1 µL of gDNA extracted from fixed cells at a final concentration of 30 pg/µL into the lysis mix. We incubated the plate at 50 °C for 1 h and 80 °C for 10 min in a PCR thermocycler. After lysis, we added 31 µL of MALBAC pre- amplification reaction mix (30 µL of pre-amp buffer + 1 µL of pre-amp enzyme mix) into each well, and centrifuged the plate at 600×g for 3 min followed by incubation at 94 °C for 3 min to denature DNA and then by 8 cycles of quasilinear amplification (20 °C for 40 s, 30 °C for 40 s, 40 °C for 30 s, 50 °C for 30 s, 60 °C for 30 s, 70 °C for 4 min, 95 °C for 20 s and 58 °C for 10 s) in a PCR thermocycler. After pre-amplification, we added 30.8 µL of MALBAC amplification mix (30 µL of amp buffer + 0.8 µL of amp enzyme mix) into each well, centrifuged the plate at 600×g for 3 min followed by incubation at 94 °C for 30 s to denature DNA and then by 14 cycles of exponential amplification (94 °C for 20 s, 58 °C for 30 s, 72 °C for 3 min) in a PCR thermocycler. We performed lysis, pre-amplification and amplification steps using reagents included in the MALBAC kit (Yikon Genomics, cat. no. Y001A). We purified the amplified material from each well separately using 1.8 vol./vol. ratio of Agencourt Ampure XP beads (Beckman Coulter, Cat. No. A63881) and measured the concentration using Qubit DNA HS kit (Thermo Fisher Scientific, Cat. No. Q32851). We spared 200 ng of DNA for the following steps. First, we sheared the samples using Covaris ME220 Focused-ultrasonicator with a target peak set at 200 base pairs (bp) and then performed library preparation using the NEBNext Ultra II DNA Library Prep Kit for Illumina (NEB, cat. no. E7645S). We checked the fragment distribution of the final libraries on a Bioanalyzer 2100 using DNA HS chip (Agilent, cat. no. 5067-4626). To reduce the MALBAC reagent volumes and therefore the cost per cell, we tested different scaled-down versions of MALBAC (sMALBAC) by reducing the reagent volumes in the lysis, pre-amplification, and amplification steps 50, 100, 200 or 500 times. To this end, we first sorted fixed or live single cells in a 384-well plate pre-filled with 5 µL of Vapor-Lock

(Qiagen, cat. no. 981611) and dispensed the scaled reagent volumes using the I.DOT nanodispenser (CELLINK) in 384-well plates. We then collected the MALBAC products manually from each well by adding 5 µL of Nuclease-Free Water to each well, and then fragmented each sample, and prepared sequencing library as described above.

**Assessment of scCUTseq sensitivity**

***Generation of Cas9-expressing TK6 cells.*** We cultured TK6 lymphoblastoid cells in RPMI-1640 medium with 5 % horse serum (Gibco, cat. no. 11510516), supplemented with 2 mM L-glutamine (Gibco, cat. no. A2916801), 100 U/ml penicillin/100 µg/ml streptomycin (Gibco, cat. no. 15140122), and 100 µM sodium-pyruvate (Gibco, cat. no. 11360070) at 37 °C and 5% $CO_2$. To establish TK6-Cas9 cells stably expressing SpCas9, we transduced the parental cells with viral particles produced using a Cas9 lentiviral expression vector (pLenti-Cas9 Blast, Addgene cat. no. 52962, kind gift from Feng Zhang). Briefly, we produced lentiviral particles in HEK293T cells by transfecting them with 4 µg of the Cas9 lentiviral expression vector and 1 µg each of lentiviral packaging plasmids, pMDLg/pRRE, pRSV-REV, pMD2.G (Addgene, cat. no. 12251, 12253, and 12259, respectively, all kind gifts from Didier Trono) using the X-tremeGENE HP DNA Transfection Reagent (Roche, cat. no. 6366244001). We pooled the virus-containing supernatant on two consecutive days from two 10 cm dishes of HEK293 transfected with the same plasmids and concentrated the supernatant 100 times using Lenti-X Concentrator (TaKaRa, cat. no. 631231). We then added the concentrated virus to 106 TK6 cells together with 8 µg/ml of Polybrene (Sigma-Aldrich, cat. no. TR-1003-G). One day after infection, we exchanged the medium and added 5 µg/ml Blasticidin to select the cells that had been successfully transfected. We prepared single-cell clones from limiting dilutions in 96-well plates and checked each clone for Cas9 expression and activity by immunoblotting and immunofluorescence microscopy using an anti-Cas9 antibody (Active Motif, cat. no. 61577), and by performing a T7 endonuclease assay following electroporation of the cells with specific gRNAs.

***Deletion induction by CRISPR/Cas9.*** To assess the sensitivity of scCUTseq, we generated a 7 Mb deletion on chr11 in the Cas9-expressing TK6 cell line described above. We targeted the *KMT2A* (hg38 chr11:118488514-118488533) and *HYLS1* (hg38 chr11:125899369-125899388) loci, which are approximately 7 Mb apart on the q-arm of chr11. To this end, we purchased guide RNAs (gRNAs) (*KMT2A*: TTTGGGTTTTAGTAGTCCAC; *HYLS1*: ATGGAAGAACTTCTACCTGA) from Dharmacon and assembled them into active complexes by adding crRNA-tracrRNA components (Dharmacon, cat. no. U-002005-5) according to the manufacturer's instructions. We electroporated the sgRNA complexes to TK6-Cas9 cells established as described above, using the Neon Transfection System (Thermo Fisher Scientific, cat. no. MPK10025) with the following program: 1350 V, 10 msec, 3 pulses. We used 106 cells with a final amount of 1.6 nmol of gRNA in each electroporation. Afterwards, we kept the cells in growing medium for two days at 37 °C and 5% CO2 to allow for the 7 Mb genomic region between the *KMT2A* and *HYLS1* loci to be deleted in a fraction of the cells.

***High-throughput DNA FISH.*** To assess the frequency of the induced 7 Mb deletion, we prepared FISH probes targeting the KMT2A-5′ (Thermo Fisher, cat. no. CTD2159M9) KMT2A-3′ (BACPAC Resources CHORI, cat. no. RP11-59N1), HYLS1-5′ (BACPAC Resources CHORI, cat. no. RP11-712D22) and HYLS-3′ (BACPAC Resources CHORI, cat. no. RP11-50B3) loci using bacterial artificial chromosomes (BACs) and performing nick translation with the Nick Translation kit (Abbott Molecular, cat. no. 7J0001) and using the following fluorescently labeled dUTPS: AlexaFluor 488-5-dUTP (Thermo Fisher Scientific, cat. no. C11397); AlexaFluor 568-5-dUTP (Thermo Fisher Scientific, cat. no. C11399); AlexaFluor 647-AHA-dUTP (Thermo Fisher Scientific, cat. no. A32763); CF405S-dUTP (Biotium, cat. no. 40004). We plated TK6-Cas9 cells onto poly-L-lysine coated 96-well glass-bottom imaging plates (PerkinElmer, cat. no. 6055308) and spun the plates at 400×g in a swing-out rotor centrifuge for 20 sec to allow for cells to get attached to the glass. Cells were fixed in 1× PBS/4% paraformaldehyde (PFA) (Thermo Fisher Scientific, cat. no. 158127) for 15 min at room temperature and washed the cells with 1× PBS three times to remove the unreacted PFA. We permeabilized the cells in 1× PBS/0.5% saponin (Sigma-Aldrich, cat. no. 47036)/0.5% Triton X-100 (Sigma-Aldrich, cat. no. T8787) for 20 min at room temperature, followed by two washes in 1× PBS and then 15 min incubation in 0.1 N HCl at room temperature. We then washed the cells in 2× SSC at room temperature and incubated in them in 2× SSC/50% Formamide (Sigma-Aldrich, cat. no. F9037) for 20 min at room temperature. We precipitated 80 ng of each labelled BAC probe by adding 3 µg of COT-1 DNA (GeneON, cat. no. 3001), 20 µg of tRNA (Invitrogen cat. no. 10702487) and 2 volumes of 100% ethanol. Probes were centrifuged at 16,000×g for 20 min at 4 °C. We resuspended the DNA pellets in 30 µL of hybridization mix containing 2× SSC/50% formamide/10% dextran sulfate (Sigma-Aldrich, cat. no. 67578)/1% Tween-20 (Sigma-Aldrich, cat. no. P9416) and added 30 µL of each probe in hybridization mix to a single well of a 96-well plate. We performed denaturation performed at 85 °C for 10 min on a slide moat (Boekel Scientific, cat. no. 10630394), followed by plate spinning at 400×g for 20 sec. We then placed the plate in humidified chamber and incubated overnight at 37 °C. The next day, we washed the wells three times with 1× SSC at 45 °C, 5 min each, followed by three consecutive washes with 0.1× SSC at 45 °C, 5 min each. Finally, we washed the wells once with 1× PBS at room temperature and stored the plate at 4 °C until imaging. We imaged the plates on the Opera Phenix high content screening confocal microscope (PerkinElmer) operated by the Harmony 4.8 software, using a 40× NA=0.8 water immersion lens (Olympus) and a 1.3 Megapixel CCD camera with pixel binning 2, corresponding to a pixel size of 299 nm. For each condition (transfection with KMT2A/HYLS1 sgRNAs or non-targeting sgRNA control), we imaged 50 fields with 11 planes in *z* per field in three technical triplicates per experiment and two biological experiments.

***Calculation of the fraction of TK6-Cas9 edited cells.*** To calculate the percentage of TK6-Cas9 cells in which the expected 7 Mb deletion between *KMT2A* and *HYLS1* had occurred, we first segmented nuclei in the DNA FISH images based on the fluorescence background signal in maximally projected images, using a custom-made pipeline built in the Harmony software. We performed spot detection of all four colored FISH spots in different channels using a built-in analysis block in the Harmony software. To determine 3D distances between spots, we calculated the Euclidean spot-to-spot distances from the position of spots in maximally projected images in *x*,*y* and the distances between the *z*-planes of their brightest pixels. We corrected the distances between the *z* planes of spots in different colors for shifts due to chromatic aberration in *z*, by determining the offset of spot detection between the channels in *z* for one genomic locus simultaneously stained with all four different colored probes. We performed all the calculations using custom made R scripts (available upon request) with text files of analyses derived from the Harmony software as input. We called cells as harboring the 7 Mb deletion when the KMT2A-3′ and HYLS1-5′ probes (see scheme in **Supplementary Fig. 2a**) were not detected, and the distance between the KMT2A-5′ and HYLS1-3′ probes was smaller than a threshold set based on their average distance in non-targeted control cells. Similarly, we called cells as harboring the deletion plus chromosome 11 arm loss, when all probes were detected only once per cell, except the KMT2A-5′, which was detected twice. Finally, we called cells as harboring the deletion plus an amplification of 3′ arm of chromosome 11, when cells had two KMT2A-5′ probes, one KMT2A-3′ probe, one HYLS1-5′ probe and three HYLS1-3′ probes.

#### Targeted DNA sequencing

To profile mutations in cancer-associated genes in each region in the P3 and P6 prostate samples, we prepared libraries from 40 ng of gDNA per region using the TruSight Oncology 500 panel (Illumina, cat. no. 20040765). This panel is 1.94 Mb in size, encompassing the full coding sequencing of 523 cancer-related genes (coding size: 1.2 Mb). We sonicated 80 ng of genomic DNA extracted from each region in P3 and P6 samples using a Covaris Focused-ultrasonicator (Covaris) and then prepared libraries and performed two rounds of hybridization-based target capture following the manufacturer's instructions. We sequenced all the libraries on Illumina NovaSeq 6000 aiming at reaching minimum 500X read depth. We processed raw data using the TruSight Oncology 500 v2.2 Local App (Illumina) to generate fastq files by aligning the reads to the human reference sequence GRCh37 (hg19). We used the same application to perform QC and somatic variant calling using the tumor-only pipeline.

**Whole genome sequencing**

To profile germline CNVs in the P3 and P6 prostate samples, we extracted gDNA from peripheral blood using the DNeasy Blood & Tissue Kit (Qiagen, cat. no. 69504). To profile bulk SCNAs in two regions of the P6 prostate sample, we extracted the gDNA from the leftover nuclei suspension after single-nucleus sorting using the DNeasy Blood & Tissue Kit as above. We prepared individual libraries from each of the gDNA samples using the NEBNext Ultra II FS DNA Library Prep Kit (New England Biolabs, cat. no. E7805L) following the manufacturer's instructions. In brief, we used 50 ng of gDNA from each sample as input. We enzymatically fragmented the gDNA at 37 °C for 30 min followed by heat inactivation at 65 °C for 30 min to achieve a target size of approximately 200 bp. Subsequently, we performed end-repair and adapter ligation in the same tube followed by purification of the fragments using a 0.8 v/v ratio of Ampure XP beads (Beckman Coulter, cat. no. A63881). We amplified adapter-ligated DNA fragments with 5 PCR cycles with barcoded primers (New England Biolabs, cat. no. E7500) and purified the PCR product with a 0.9 v/v ratio of Ampure XP beads. We sequenced the peripheral blood gDNA libraries on NextSeq 2000 (Illumina) with pair-end mode using the NextSeq 1000/2000 P3 Reagents (300 Cycles) kit (Illumina, cat. no. 20040561), while the tissue block libraries were sequenced with single-end mode using the NextSeq 1000/2000 P2 Reagents (100 Cycles) kit (Illumina, cat. no. 20046811). See **Supplementary Table 9** for a summary of sequencing statistics.

#### 3. Supplementary Tables

Because of their large size, all Supplementary Tables are provided as separate Excel files.

**Supplementary Table 1.** Pathological characteristics of the prostate samples used in this study.

**Supplementary Table 2.** Concentration and integrity of the RNA extracted from the regions in prostate samples P2 and P5 profiled by scCUTseq.

**Supplementary Table 3.** List of metrics describing manually annotated copy number profiles used to train a random forest classifier.

**Supplementary Table 4.** Characteristics of the donors of the brain and skeletal muscle samples profiled by scCUTseq in this study.

**Supplementary Table 5.** Sequence of the oligos composing the DNA FISH probes used for scCUTseq validation.

**Supplementary Table 6.** Frequency of amplification or deletion of COSMIC genes in TRR/FER-specific and in the five most localized pseudo-diploid subclones in prostate samples P2 and P5.

**Supplementary Table 7.** List of genes mutated in prostate samples P2 and P5.

**Supplementary Table 8.** Sequence of the CUTseq oligonucleotide adapters used in this study.

**Supplementary Table 9.** Summary of sequencing runs.

**Supplementary Table 10.** List of reagents for scCUTseq, standard MALBAC, and ACT and relative costs.

#### 4. Supplementary Notes

##### 1. Cumulative cost analysis

To assess the cost-effectiveness of scCUTseq, we first compared it to the standard MALBAC method[6], which in scCUTseq is used to perform whole-genome amplification prior to barcoding genomic DNA (gDNA) in individual cells (see **Fig. 1a**). In scCUTseq, the volume of MALBAC reagents used in the lysis, pre-amplification and amplification steps is 200-fold lower compared to standard MALBAC. Furthermore, while in the latter a single sequencing library needs to be generated from a whole genome amplified cell, in scCUTseq multiple cells (typically, 96 or 384, depending on the number of CUTseq adapters available) are pooled into the same library after barcoding gDNA in each individual cell. A list of reagent costs (based on prices as of June 2021) used to make the cost comparison is provided in **Supplementary Table 10**. Note that sequencing costs are excluded, since they only depend on the number of single cells sequenced and target sequencing depth. As shown in the **Supplementary Notes Figure 1** below, the cumulative cost grows at a much faster rate for standard MALBAC compared to scCUTseq and needs to be plotted on logarithmic scale for the difference between the two curves to be visible. This is expected from the fact that in scCUTseq 200-fold less MALBAC reagents are used, and 384 cells are pooled into the same library, drastically reducing the cost per cell. Assuming to sequence 10,000 cells as we did in this study, the reagent costs for making libraries would be 33.6-fold higher using standard MALBAC compared to scCUTseq (1,123,221 USD *vs.* 33,335 USD, respectively), highlighting the cost-effectiveness of scCUTseq.

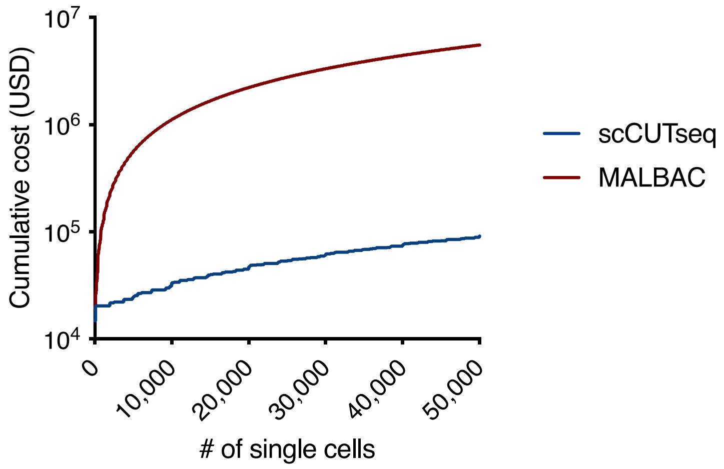


**Supplementary Notes Figure 1.** Plot showing how the cost for preparing libraries for single-cell copy number profiling grows with the number of single cells processed using standard

MALBAC (red) *vs.* scCUTseq (blue). Note that the y-axis in on logarithmic scale to be able to distinguish between the two curves.

Next, we performed a side-by-side comparison with acoustic cell tagmentation (ACT)[7], a scDNA-seq method that, similarly to scCUTseq, also uses a nanodispensing device (Echo 525, Beckman Coulter) to reduce the volume of reagents delivered to each single cell in 96- or 384-well plates. However, unlike scCUTseq, ACT uses a scaled-down version of the Illumina's Nextera library preparation kit (25-fold less reagent volumes per cell compared to the recommended volumes per sample in the Nextera kit) to index individual cells with the Tn5 transposase, followed by 18–16 polymerase chain reaction (PCR) cycles before pooling multiple cells together for sequencing. To compare the two methods, we computed the cumulative costs of preparing ready-to-sequence libraries for 50,000 cells using scCUTseq *vs.* ACT, assuming to use 384 CUTseq adapters (as we did in this study) and purchasing all the 384 indexes available in the Nextera kit. All the reagent costs used for this analysis are listed in **Supplementary Table 10**. We did not include sequencing costs in our analysis, as these would be the same for scCUTseq and ACT, for any given target depth per cell. As shown in the **Supplementary Notes Figure 2** below, up to 7,296 single cells processed, ACT is more cost-effective compared to scCUTseq. Above 7,296 cells, scCUTseq becomes progressively more cost-effective. For 10,000 cells (approximately the number of single cells from prostate samples that we sequenced in this study), scCUTseq is ~10% less costly than ACT, while for 50,000 cells the difference would increase to ~77%.

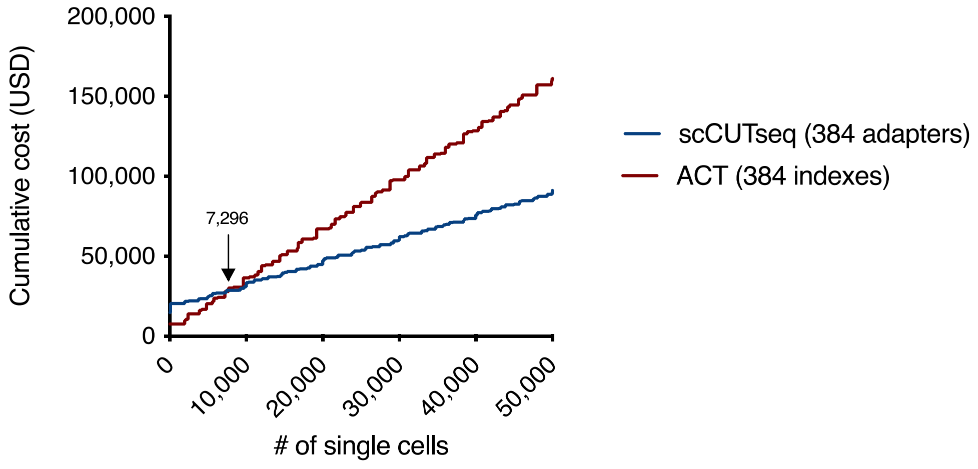


**Supplementary Notes Figure 2.** Plot showing how the cost for preparing libraries for single-cell copy number profiling grows with the number of single cells, using scCUTseq *vs.* ACT.

The higher cost for scCUTseq below 7,296 cells can be explained by the need to initially purchase more reagents compared to ACT. However, as shown in **Supplementary Table 10**, many of these reagents are then sufficient to process thousands or even millions (as in the case of CUTseq oligo adapters) of cells, thus making scCUTseq less costly than ACT above 7,296 cells.
